# TCR-BERT: learning the grammar of T-cell receptors for flexible antigen-xbinding analyses

**DOI:** 10.1101/2021.11.18.469186

**Authors:** Kevin Wu, Kathryn E. Yost, Bence Daniel, Julia A. Belk, Yu Xia, Takeshi Egawa, Ansuman Satpathy, Howard Y. Chang, James Zou

**Affiliations:** Department of Computer Science, Stanford University, Stanford, CA 94305; Center for Personal Dynamic Regulomes, Stanford University, Stanford, CA 94305; Department of Pathology, Stanford University School of Medicine, Stanford, CA 94305; Department of Pathology and Immunology, Washington University School of Medicine, St. Louis, MO 63110; Howard Hughes Medical Institute, Stanford University, Stanford, CA 94305; Department of Biomedical Data Science, Stanford University School of Medicine, Stanford, CA 94305

## Abstract

The T-cell receptor (TCR) allows T-cells to recognize and respond to antigens presented by infected and diseased cells. However, due to TCRs’ staggering diversity and the complex binding dynamics underlying TCR antigen recognition, it is challenging to predict which antigens a given TCR may bind to. Here, we present TCR-BERT, a deep learning model that applies self-supervised transfer learning to this problem. TCR-BERT leverages unlabeled TCR sequences to learn a general, versatile representation of TCR sequences, enabling numerous downstream applications. We demonstrate that TCR-BERT can be used to build state-of-the-art TCR-antigen binding predictors with improved generalizability compared to prior methods. TCR-BERT simultaneously facilitates clustering sequences likely to share antigen specificities. It also facilitates computational approaches to challenging, unsolved problems such as designing novel TCR sequences with engineered binding affinities. Importantly, TCR-BERT enables all these advances by focusing on residues with known biological significance. TCR-BERT can be a useful tool for T-cell scientists, enabling greater understanding and more diverse applications, and provides a conceptual framework for leveraging unlabeled data to improve machine learning on biological sequences.

## Introduction

T cells are a central component of the adaptive immune system ^1^. Mature T cells continuously monitor their surroundings – in the blood and in tissues – for signs of foreign invaders or diseased cells and help activate other immune defenses upon recognition. In the event of a viral infection such as with HIV or SARS-CoV-2, for example, infected cells present viral antigens (i.e., short viral peptide chains) on the peptide-major histocompatibility complex (pMHC), which are then recognized by T cells ^2^. In cancer, fragments of mutated intracellular proteins are similarly presented on the MHC by cancerous cells and detected by the immune system as neoantigens. T-cell mediated anti-tumor immunity can be therapeutically harnessed in several ways, including immune checkpoint blockade therapies which show clinical benefit in a variety of cancer types ^3^. Healthy cells similarly present antigens identifying themselves as non-threatening, but aberrant recognition of these self-antigens as invaders – a phenomenon frequently known as autoreactivity – can cause autoimmune disorders like multiple sclerosis ^4 5^ and type 1 diabetes ^6 7^. In each of these settings, understanding the antigen(s) recognized by T cells is key to understanding the underlying drivers and progression of each malignancy and for developing effective treatments ^8 9 10 11 12 13^.

Despite the importance and therapeutic potential of T cells, it is challenging to predict their antigen recognition behavior, which is mediated via the T-cell receptor (TCR). The TCR itself is a dimeric protein with two hypervariable chains – typically α and β chains, encoded by the *TRA* and *TRB* genes – that jointly bind to and interact with the pMHC-antigen complex ^14^. This biophysical interaction is believed to be primary determinant of antigen specificity ^15 16^. These TRA and TRB sequences are specified through recombination of the variable (V), diversity (D), and junction (J) gene segments, as well as through random insertions and deletions. This stochastic process generates a staggering diversity of different TCR sequences – often estimated to be on the order of (hundreds of) millions for a healthy human individual ^17 18 19^. This diversity crucially lends the immune system its ability to recognize a vast array of antigens, but also makes precisely understanding and predicting TCR-antigen specificity difficult. This challenge is compounded by observations that a single TCR often recognizes multiple antigens ^20 21 22^, a phenomenon known as cross-reactivity, and conversely, that any given antigen may be recognized by many TCRs with varying affinities ^23^.

Recently, the growing popularity and accessibility of sequencing technologies has enabled high-throughput profiling of TCR sequences ^24 25^, which has in turn empowered numerous computational methods studying and predicting TCR-antigen binding. Conventional approaches like GLIPH ^15 26^, TCRMatch ^27^, and TCRdist ^28^ typically rely on sequence motif comparisons and manually engineered heuristics to predict which TCRs are likely to share common antigen binding partners. More recently, researchers have applied various machine learning methods to predicting TCR-antigen binding. Examples include DeepTCR (variational convolutional autoencoder) ^29^, SETE (gradient boosting decision tree) ^30^, and TcellMatch (multiple deep learning architectures) ^16^. While the specific modelling approaches of these tools differ, they share the same core approach of taking a set of TCR sequences (and optionally VDJ gene usage) that are individually annotated, or labeled, with antigen binding information and using these labelled examples to train a classifier predicting binding. This approach of using only labelled data is known as supervised learning. However, many of the TCR sequences that scientists have collected do not have corresponding antigen labels required for such supervised approaches. Furthermore, it is likely that existing labels are incomplete owing to cross-reactivity. Thus, there is a unique opportunity to develop new approaches that leverage this wealth of unlabeled data in concert with labelled examples to improve TCR models’ robustness and generalizability.

This challenge of using unlabeled data to build models is a burgeoning field of study within the broader machine learning community. A notable recent innovation in this domain is the Bidirectional Encoder Representations from Transformers (BERT) architecture, originally developed for natural language processing tasks ^31^. Colloquially, BERT is a highly expressive model trained to understand the grammatical structure of languages using large, heterogeneous collections of unlabeled sequences – a process more formally known as pre-training. This general understanding then serves as a robust starting point for quickly and effectively targeting more specific tasks like predicting whether text exhibits positive or negative sentiment. While BERT was originally designed to model sequences of words comprising human language, BERT and its related architectures have recently been successfully repurposed for modelling biological sequences like DNA ^32^ and proteins ^33 34 35^.

Inspired by the challenges of TCR analysis and the success of these machine learning approaches, we present TCR-BERT, a modified BERT architecture trained specifically on TCR amino acid sequences. TCR-BERT explicitly leverages unlabeled TCR sequences to achieve state-of-the-art performance on a wide variety of downstream tasks and applications in TCR analysis. First, TCR-BERT enables class-leading prediction of TCR specificity. These predictors stand out in their flexibility and generalizability, particularly when trained and evaluated across different patients. We also show that TCR-BERT’s learned embedding can be used to identify sets of TCRs likely to share antigen specificities, and that TCR-BERT provides tangible improvements over prior tools with similar aims. We then demonstrate that in enabling these advances, TCR-BERT learns to focus on residues with well-established biophysical importance despite being given only raw, unannotated amino acid sequences. We also present a proof-of-concept showing how TCR-BERT can serve as foundation for future technologies, such as by facilitating *in silico* design of novel TCR sequences with predetermined, desirable antigen binding characteristics. Overall, we believe TCR-BERT is an important step towards a generalizable computational understanding of TCRs that enables advances in computational biology, biological understanding, and clinical applications.

## Results

### TCR-BERT leverages large, unannotated datasets to learn representations of TCRs

TCR-BERT is built upon a modified BERT architecture (see Methods for details) that takes TCR amino acid sequences as input (e.g., CASRPDGRETQYF). We pre-train TCR-BERT to capture the language of TCR sequences by optimizing two objectives sequentially. First, we use unlabeled TCR sequences to learn the underlying grammar of the naturally occurring TCR sequence space. We randomly hide 15% of the residues in each sequence and train TCR-BERT to fill in these hidden amino acids based on surrounding residues. Critically, this masked amino acid (MAA) pre-training does not require knowledge of the antigen binding partners of each TCR sequence. MAA pre-training is performed using a set of 88,403 predominantly human TRA and TRB sequences drawn from the public VDJdb ^36^ and PIRD ^37^ datasets (Figure 1). These sequences span a wide range of known and unknown antigen specificities, as well as other phenotypes like HLA alleles.

**Figure 1:**
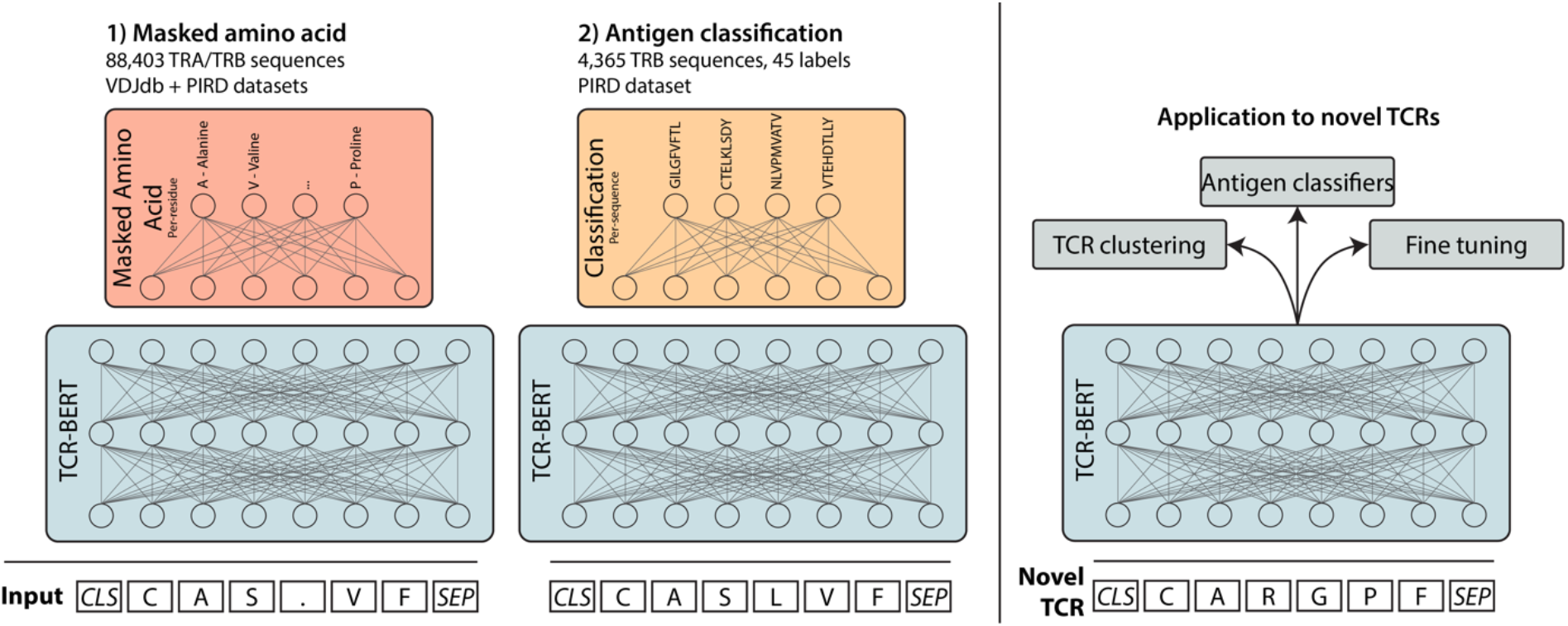
TCR-BERT leverages self-supervised pretraining to model TCRs. TCR-BERT takes a T-cell receptor amino acid sequence and generates a continuous embedding that can be used for downstream tasks. To pre-train TCR-BERT, we first perform masked amino acid prediction, training TCR-BERT to predict a masked or hidden amino acid (“.” in the input) based on surrounding amino acids, thus learning the “grammatical” structure of naturally occurring TCRs. This is done over a large corpus of TRA and TRB sequences with no MHC or HLA restrictions and crucially does not require knowledge of antigen binding affinities (left panel). Next, we take this model and further train it to predict, across a set of 45 antigen labels, the antigen that a given TRB amino acid sequence binds to (center panel). After checking pre-training’s efficacy (Supplementary Figure 1) and selecting an optimal representation layer for downstream tasks (Supplementary Figure 2), TCR-BERT can be used for a variety of TCR analyses, including predicting antigen binding and clustering TCRs (right panel).

After MAA prediction, we leverage the fact that some TCRs are labelled with known antigen binding to train TCR-BERT to predict antigen specificity given TCR sequence. We use 4,365 human TRB sequences, each binding to one of 45 antigen labels derived from the PIRD dataset (Figure 1, see Methods for additional details). Although this antigen classification pre-training step uses relatively few examples, it provides substantial benefits to TCR-BERT (see following results section). However, since antigen classification pre-training only considers TRB sequences (as there are insufficient labelled TRA sequences), TCR-BERT is better suited for processing TRB sequences after this pre-training step. For applications studying both TRA and TRB sequences, we recommend using the version of TCR-BERT pretrained only on MAA.

We performed several evaluations to check the quality and generalizability of TCR-BERT’s pre-training using a new dataset of (n=17,702) TRA/TRB sequence pairs from mice infected with lymphocytic choriomeningitis virus (LCMV) clone 13 antigen GP33, a model of chronic viral infection (see Methods for additional details). This dataset is fully external to model training and is murine whereas 97% of TCR-BERT’s pre-training data comes from humans. Nonetheless, TCR-BERT accurately predicts masked amino acids for both TRA and TRB sequences (Supplementary Figure 1A) after MAA pre-training, suggesting that the model has learned a generalizable TCR grammar. After classification pre-training, we visualize the amino acid embedding vectors that TCR-BERT has learned and observe expected separation by biochemical properties (Supplementary Figure 1B). We additionally take the murine LCMV dataset, use TCR-BERT to generate an embedding vector for each TRB sequence, and visualize the embeddings using UMAP ^38^ (Supplementary Figure 1C). We observe that TCR-BERT’s sequence embedding reflects sequence similarity (Supplementary Figure 1C, 1D). As sequence similarity is a known predictor of similar antigen binding affinities ^15 39^, this suggests that TCR-BERT’s sequence embeddings capture meaningful structures and relationships within the TCR sequence space and are likely a powerful basis for predicting antigen affinity – a topic that we explore further in the following section.

### TCR-BERT enables state-of-the-art antigen specificity classifiers

After pre-training, we use TCR-BERT as a basis for building classifiers to predict whether a given TCR sequence can bind to a specific antigen. For this, we treat TCR-BERT as a fixed black box for generating embedding vectors for TCR sequences (Supplementary Figure 2). We project the TCR embeddings onto its top 50 principal components (PCs) and train a support vector machine (SVM) to predict whether TCRs bind to a specific antigen. We refer to this classification module as PCA-SVM (see Methods for additional details).

To evaluate the efficacy of applying PCA-SVM to TCR-BERT’s embedding, we perform antigen cross validation. For a subset (n=26) of the 44 antigens used in antigen classification pre-training with at least 20 known binding TRBs, we repeat our second antigen classification pre-training step excluding that antigen and its associated TRBs. We then use the resulting model to embed and classify the held-out TRBs against a negative set of human TRBs of undetermined affinity using PCA-SVM. This negative set is randomly sampled from the TCRdb database ^40^, which is not seen in pre-training, at a ratio of 5 negatives to each binding TRB (see Methods for additional details). Using a random 70/30 train/test split, we observe a test set area under the precision-recall curve (AUPRC) of 0.847 averaged across the 26 antigens, compared to an AUPRC of 0.167 expected for a random classifier.

We similarly applied antigen cross-validation to several other antigen-TCR classifiers to contextualize TCR-BERT’s performance. As a baseline, we evaluated a supervised convolutional network (ConvNet) deep learning model classifying antigen binding/non-binding given a TRB sequence (see Methods for details). We train this ConvNet from scratch (i.e., without pre-training) to classify each of the 26 antigens’ TRBs against the negative background described above. For 25 of the 26 antigens, TCR-BERT exhibits improved AUPRC compared to this supervised ConvNet (Figure 2A, *p* = 4.67 × 10^−6^, Wilcoxon test). We find similar improvements when comparing TCR-BERT against other supervised methods like SETE ^30^ (Supplementary Figure 3A). These results suggest that pre-training strategies provide tangible performance improvements over supervised models trained from scratch.

**Figure 2:**
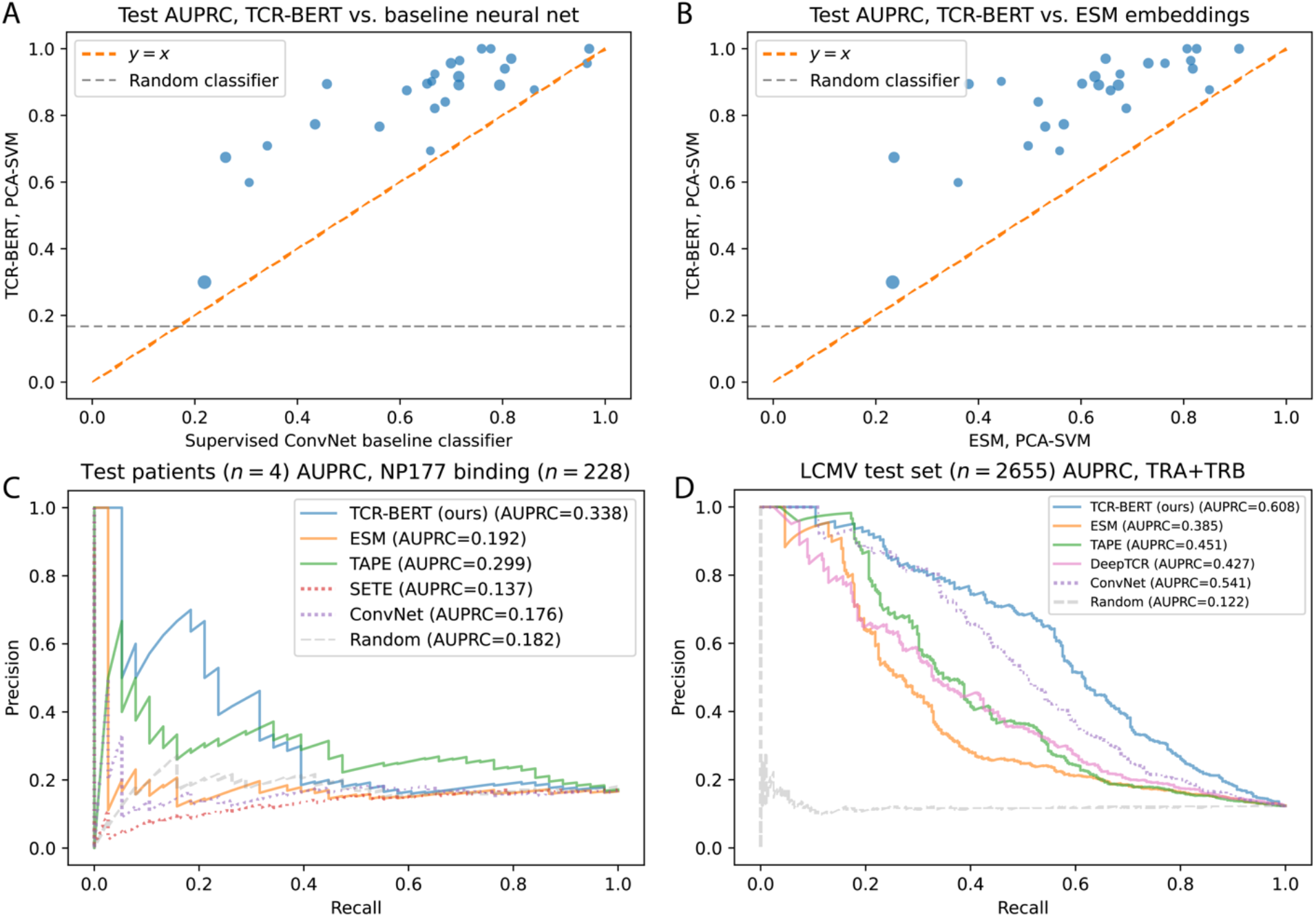
TCR-BERT can be used to build state-of-the-art classifiers predicting antigen binding. (A) Antigen cross validation comparing PCA-SVM on TCR-BERT embeddings (y-axis) to a baseline supervised convolutional neural network (ConvNet, x-axis). Each point represents test AUPRC classifying a single antigen using each of the two methods. Larger points indicate antigens with more training examples available (log-scaled). TCR-BERT delivers improved performance (i.e., above the orange line indicating equal performance) in 25/26 instances. We observe similar improvements when comparing against other supervised methods (Supplementary Figure 3A). (B) Antigen cross validation applying PCA-SVM to our TCR-BERT model (y-axis) compared to PCA-SVM on a similar language model targeting general amino acid sequences, ESM (x-axis). Each point represents AUPRC classifying a single antigen using PCA-SVM. In all cases, TCR-BERT’s embedding enables substantially improved classification performance. This holds for other large amino acid language models like TAPE as well (Supplementary Figure 3B), which suggests that TCR-BERT’s specialized pre-training is critical to achieving good performance on these hypervariable TCR chains. (C) We evaluate various antigen binding prediction methods’ ability to generalize across different patients. We train classifiers to predict TRB binding to the human NP177 antigen using a single patient’s data and evaluate these models on 4 test patients. Models that leverage pre-training on large datasets are shown in solid lines whereas supervised models are shown in dotted lines. TCR-BERT and TAPE are the only methods that predict TCR-antigen specificity generalizably across patients, with TCR-BERT providing the best performance. On the other hand, both supervised methods perform worse than a random classifier, suggesting overfitting. (D) Using a murine dataset profiling GP33 binding, we evaluate various models’ antigen binding predictions when given TRA/TRB sequence pairs. We fine-tune TCR-BERT to achieve class-leading performance (see also Supplementary Figure 4), as it did for previous TRB-only antigen binding prediction problems. As in panel (C), pre-trained models are shown using solid lines, and supervised models are shown in dotted lines.

As another benchmark, we apply the same PCA-SVM classification module, but rather than using TCR-BERT’s embeddings, we use embeddings generated by ESM, a substantially larger transformer model trained on general amino acid sequences ^33^. We find that for all 26 antigens, TCR-BERT’s embedding yields improved AUPRC compared to ESM’s embedding (Figure 2B, *p* = 4.15 × 10^−6^, Wilcoxon test). This result holds for other general-purpose amino acid transformer models like TAPE ^34^ (Supplementary Figure 3B). In these comparisons, the classifier module is held constant while the embedding representation varies, thus demonstrating that TCR-BERT’s TCR-specific pre-training results in a more powerful TCR embedding representation compared to general-purpose protein language models.

Antigen cross-validation also allows us to investigate the impact of each of our two pre-training steps on the quality of TCR-BERT’s embeddings. We compare PCA-SVM’s performance with an embedding trained using only MAA, as well as an embedding trained using only antigen classification (holding out antigens as necessary). We find that for both these pre-training ablations, AUPRC classifying unseen antigens is significantly reduced (Supplementary Figures 3C, 3D). This highlights the importance of both our pre-training steps in learning an embedding that generalizes to unseen antigens and their associated TRBs.

We additionally sought to understand how well these performance gains would generalize not just across random data splits, but across patients. As each individual’s immune system independently generates TCR sequences, cross-patient generalization better captures how such antigen classifiers might be applied in a clinical setting. We focus on (n=214) human TRB sequences binding to the NP177 influenza A viral antigen (antigen sequence LPRRSGAAGA) ^15^, which was not seen during pretraining. These TRBs are primarily derived from a single patient (n=176), whose data we use for training, with the remaining (n=38) TRBs derived from 4 other patients, whose data we use for testing. Since this dataset only provides positive examples, we again add TCRdb human TRBs as background negatives at a ratio of 5:1. Among all evaluated models (PCA-SVM on TCR-BERT, PCA-SVM on ESM, PCA-SVM on TAPE, ConvNet, and SETE), only PCA-SVM on TCR-BERT and PCA-SVM on TAPE meaningfully generalize to test patients, with TCR-BERT providing the best AUPRC on test patients (Figure 2C). Notably, both models with nontrivial generalization were pre-trained on large corpuses of unlabeled data. This strongly suggests that pre-training is critical for robust generalizability. Neither supervised approach (ConvNet and SETE) performs better than random on test patients, suggesting that using only direct supervision leads to overfitting to the training patient’s TRBs.

Finally, we evaluate classification performance on the murine LCMV GP33 dataset. This dataset uniquely provides antigen binding annotations for many (n=17,702) TRA/TRB pairs. To leverage the TRA/TRB pairing information, we fine-tuned two separate copies of TCR-BERT, both initialized using weights from MAA pretraining, to embed TRA and TRB sequences, respectively. These two embeddings are concatenated and passed through a single-layer fully-connected classification head (see Methods for additional details). We compare this method to several approaches. As a baseline, we train a ConvNet with two convolutional “arms” corresponding to the TRA and TRB, respectively. We evaluate TAPE and ESM by applying PCA-SVM on their concatenated representations of TRA and TRB. We compare our method against DeepTCR, a variational autoencoder embedding TRA/TRB pairs ^29^, which we train on other murine TRA/TRB pairs before embedding our GP33 TCRs (see Methods for details). We then apply a PCA-SVM classifier on these embeddings. Compared to all methods, TCR-BERT achieves best-in-class performance predicting binding to GP33 with an AUPRC of 0.608 (Figure 2D, Supplementary Figure 4). This strong performance is largely a consequence of TCR-BERT’s pre-training: similarly training an architecturally identical but randomly initialized classifier achieves a substantially lower test AUPRC of only 0.462.

Overall, our results demonstrate that TCR-BERT provides a strong, versatile foundation for building antigen classifiers for TCR sequences. This can be achieved via treating TCR-BERT as a black box for generating sequence embeddings, which is typically optimal for smaller datasets, or by using larger datasets to fine-tune TCR-BERT, as we performed for LCMV GP33. Furthermore, we demonstrate that TCR-BERT can be flexibly used to predict binding for TRB sequences alone, or for TRA/TRB sequence pairs.

### TCR-BERT facilitates unsupervised, explorative analyses of TCRs

In many cases, researchers may not know *a priori* which specific antigen(s) to predict TCR affinity for, necessitating more exploratory analyses of TCR sequence data. One common task is to identify groups of TCRs likely to share antigen binding properties, which can help identify motifs and often provides a more succinct representation of the TCR repertoire than analyzing each individual clone. TCR-BERT facilitates such analyses via its TCR sequence embeddings. To illustrate this, we embed (n=2,067) human TRB sequences ^15^ using TCR-BERT (pre-trained on MAA and classification) and visualize each TRB using UMAP coloring by known antigen (Figure 3A). We see that TCR-BERT’s embedding highlights several distinct specificity groups. For example, the group of TRBs corresponding to the BMLF1 Epstein Barr virus (EBV) antigen (antigen sequence GLCTLVAML) in the upper region of the plot corresponds to the illustrated motif (Figure 3A, top right). This motif matches one of the most conserved public (i.e., shared between multiple individuals) EBV TRBs ^41^, which may contribute to its separation from other TRB sequences in this set.

**Figure 3:**
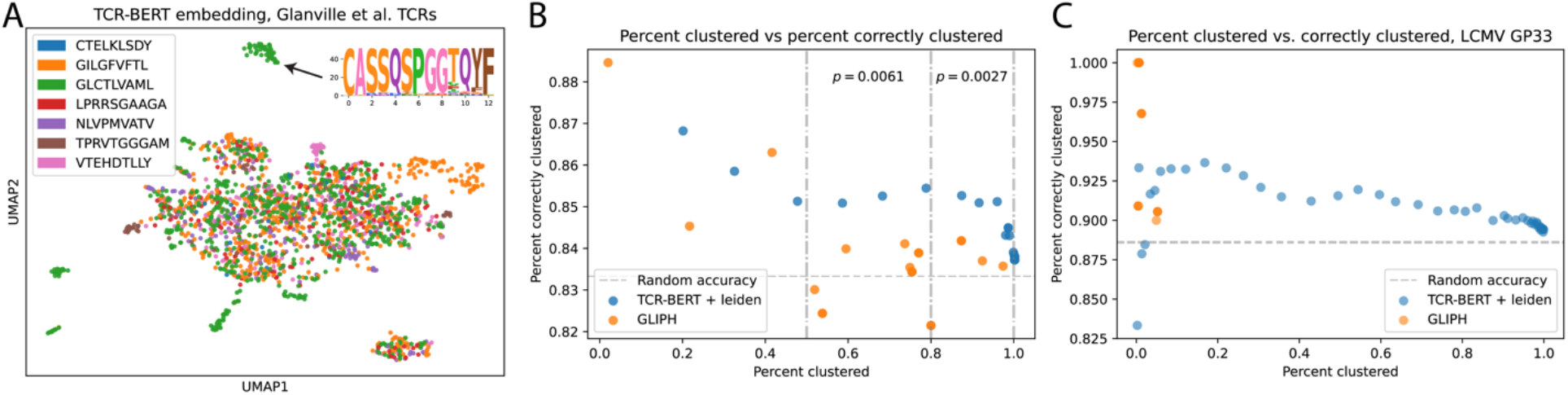
TCR-BERT’s embedding enables clustering analyses of patient TCR sequences. (A) We use TCR-BERT to embed and visualize 2,067 human TRB sequences. Each point represents a TRB sequence, colored by its known antigen binding partner. Several groups of TRBs stand out. For example, the top clusters consistently binds the GLCTLVAML antigen (green) and corresponds to the displayed consensus motif, which is highly conserved across individuals. (B) To quantify the utility of TCR-BERT’s embedding for clustering, we study (n=217) TRB sequences binding the NP177 antigen along with a negative background of randomly selected human TRBs (n=1070). We evaluate several clustering resolutions for TCR-BERT (blue) and the popular existing method GLIPH (orange), plotting each method’s trade-off between percent clustered and percent correctly clustered. TCR-BERT consistently provides comparable or improved clustering. Within each range of percent clustered (delineated by dashed vertical lines), TCR-BERT achieves improved clustering accuracy (indicated p-values, Mann-Whitney test, see Supplementary Figures 5A, 5B for additional details). (C) These improvements are consistent across different datasets, such as the LCMV GP33 murine dataset, subsampled to 2,443 TRBs, shown here. TCR-BERT (blue) provides more consistent clustering performance across a range of percent clustered values when compared to GLIPH (orange), which cannot cluster more than a handful of sequences. Note that variability in correctness at low percent clustered is the result of only a few TRBs being correctly or incorrectly classified. Plots comparing performance at detailed cutoffs are shown in Supplementary Figures 5C, 5D. These improvements are achieved with drastically improved runtime scalability (Supplementary Figure 6).

In addition to examining these embeddings visually, we can also algorithmically group these TCRs using a clustering algorithm like Leiden ^42^. We evaluate this approach using NP177-binding TRB sequences intermixed with randomly sampled endogenous human TRBs. We previously used this dataset to evaluate classification, but here we are instead interested in quantifying how well TCR-BERT’s embedding clustering separates antigen versus background TCRs. Ideally, every cluster of TRBs should homogeneously consist of either antigen-binding or background sequences; this is measured by the percent correctly clustered metric, originally proposed by the authors of GLIPH ^15^. Additionally, each TRB should be clustered with other TRBs, as assigning each TRB its own group would trivially achieve perfect accuracy; this is captured by the percent clustered metric, also proposed by the authors of GLIPH. We examine several clustering resolutions for TCR-BERT’s (pretrained on MAA and classification) NP177/background embedding, which yields a smooth tradeoff between percent clustered and percent correctly clustered (Figure 3B, blue). We use GLIPH ^15^, a well-known methodology for grouping TCRs using sequence similarity heuristics, to similarly generate TCR groups of varying granularities for these same sequences and similarly plot its relationship between percent clustered and percent correctly clustered (Figure 3B, orange). Comparing these two methods, clustering TCR-BERT’s embedding consistently produces improved results. For a target “percent clustered” of 50-80% of our input sequences, TCR-BERT’s clustering yields significantly higher clustering correctness within that interval than GLIPH (p=0.0061, Mann-Whitney test). This similarly holds for a target “percent clustered” of 80-100% (p=0.0027, Mann-Whitney test). Furthermore, in several cases where a majority of TRBs have been clustered, GLIPH’s percent correctly clustered is worse than random chance (83.3%, as 5/6 of the data are negative background examples). More detailed plots including specific parameters used for each method are included in Supplementary Figures 5A and 5B. We also note that TCR-BERT enables these improvements while achieving faster runtime compared to GLIPH (Supplementary Figure 6).

We repeated the TCR clustering comparison using a subset (n=2,443) of the murine LCMV GP33 TRBs as GLIPH cannot scale to feasibly run on the full dataset. Although this dataset provides paired TRA/TRB sequences, we focus on TRBs alone for compatibility with GLIPH. TCR-BERT (Figure 3C, blue) again provides improved clustering compared to GLIPH (Figure 3C, orange), which never clusters more than 5.2% of sequences across the (n=18) cutoff values we evaluated. TCR-BERT (pretrained on MAA and classification, Figure 3C blue) provides a predictable tradeoff between percentage clustered and percentage correctly clustered. There are a few instances of erratic correctness values when TCR-BERT clusters 2% or fewer sequences, as a handful of mistakes can have an outsized effect on correctness when clustering so few sequences. Full details, including hyperparameters used, are included in Supplementary Figures 5C and 5D. We additionally explored leveraging TCR-BERT’s TRA embeddings for clustering, but this does not improve clustering metrics for this LCMV dataset. These two experiments across two different datasets suggest that TCR-BERT’s embedding can be leveraged to perform state-of-the-art exploratory visualization and clustering analyses of TRB sequences.

### TCR-BERT focuses on biologically relevant residues

Thus far, we have shown that TCR-BERT is a versatile model that can be used for a variety of TCR analyses from predicting antigen specificity to exploratory clustering of TCRs. To better understand how TCR-BERT achieves these advances, we study the amino acid residues highlighted via TCR-BERT’s transformer attention mechanism. At a high level, attentions ^43^ dynamically capture how much a given token’s representation (here, our classification embedding) is influenced by representations of other tokens (here, each amino acid in the TCR).

We examine the TCR-BERT variant pre-trained on MAA and fine-tuned to predict LCMV GP33 binding from TRA/TRB pairs. Examining test set TRA/TRB pairs of uniform length (12 and 14 residues for TRA and TRB, respectively), we average the fine-tuned TCR-BERT TRA/TRB submodules’ per-residue attentions across these examples (n=157 spanning 36 binding and 121 non-binding examples) which yields two matrices of TRA and TRB attentions, respectively (Figure 4A). At a per-residue scale, TCR-BERT’s attentions tend to be concentrated towards the central region of both the TRA and TRB. This aligns well with prior works describing a functional “hot spot” around central residues that enables fine discrimination of different antigens ^44^.

**Figure 4:**
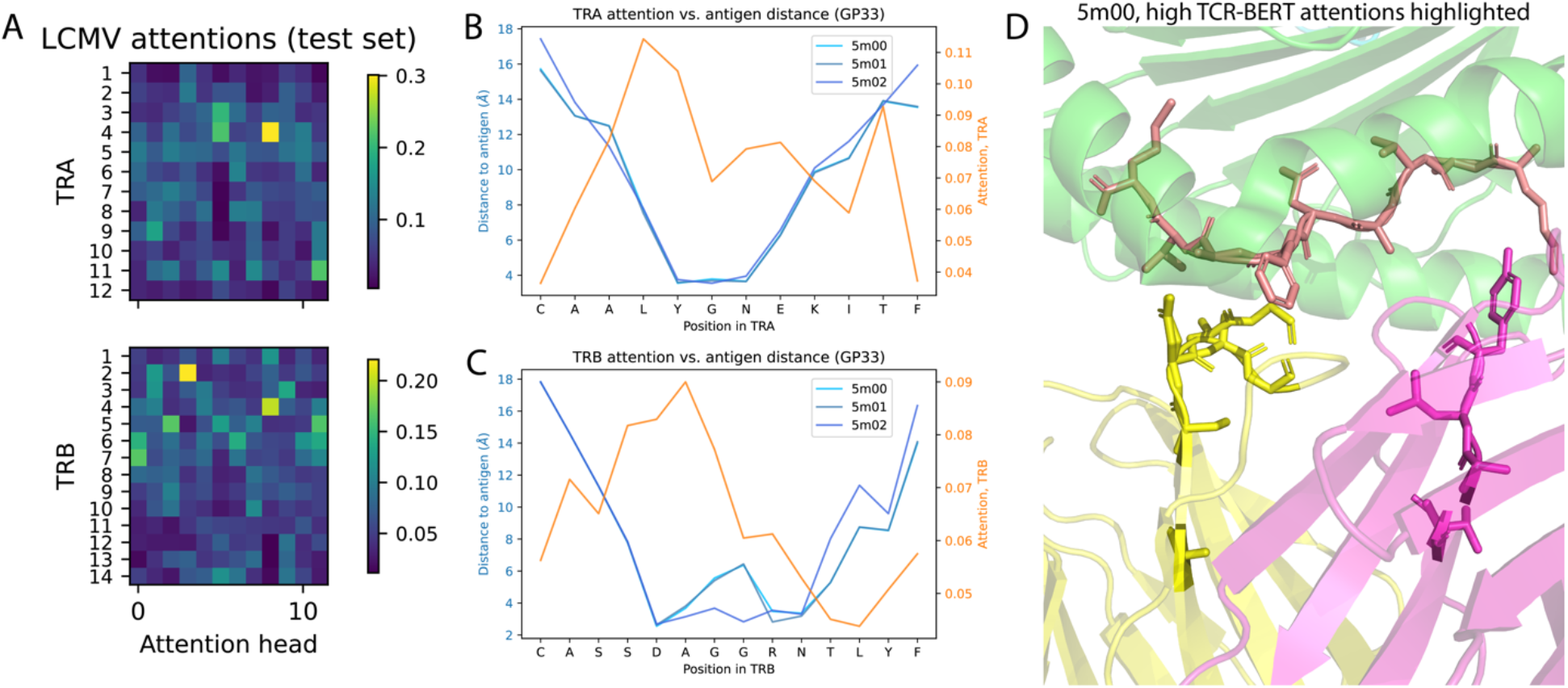
TCR-BERT’s attentions reveal biologically meaningful learned patterns. (A) Heatmaps visualizing the attentions that TCR-BERT has learned for predicting LCMV GP33 binding after fine-tuning, averaged across test set TRA and TRB sequences of fixed length. The vertical axis indicates positions in the TRA (top) and TRB (bottom) sequence, and the horizontal axis illustrates each of the 12 attention heads within TCR-BERT. Attentions tend to be concentrated to the center of the TCRs, which corresponds to prior literature. (B) We relate these averaged attentions to biophysical structures of TCR-antigen binding using three empirical PDB structures (5m00, 5m01, and 5m02) profiling a similar GP33 system. Blue lines indicate, for each residue in the TRA, the minimum distance to the antigen in each experimental structure. The orange line indicates TCR-BERT attention for those same residues. TCR-BERT pays the most attention to residues closest to the antigen, i.e., residues that are also most likely to contact the antigen. (C) The same is true for the TRB sequence. (D) 3D structure showing the MHC (green), modified GP33 antigen (salmon), and TRA (pink) and TRB (yellow). Side chains are shown and highlighted for the antigen and the TCR residues receiving the top 33^rd^ percentile of model attentions on average. The residues that TCR-BERT pays attention to are frequently in direct contact with the presented antigen; this is especially true of the yellow TRB chain, which is more proximal to the antigen. See Supplementary Figure 7 for additional views.

TCR-BERT’s attentions can also be interpreted in the context of the structural interface between the TCR and pMHC. Intuitively, TCR residues physically proximal to the antigen should play a key role in determining specificity. We sought to validate this using three publicly available experimental structures profiling a similar LCMV GP33 system (PDB structures 5m00, 5m01, and 5m02, see Methods for additional details). We calculate, for each residue in the PDB structures’ TRA and TRB sequences, the minimum Euclidean distance to any residue in the antigen (Figure 4B and 4C, blue lines). Comparing this antigen-distance to TCR-BERT’s averaged attention per position (Figure 4B and 4C, orange line), we find that attention is anti-correlated with distance to the antigen for both TRA and TRB chains. Antigens in the bottom quartile of distances relative to the antigen in each chain correspond to significantly higher attentions across these test examples (*p* = 4.6 × 10^−16^ for TRA, *p* = 5.6 × 10^−21^ for TRB, Mann-Whitney test), indicating that TCR-BERT pays the most attention to TCR residues most physically proximal to the antigen – a pattern concordant with known principles of TCR specificity. We illustrate this relationship between attention and physical proximity in a render of the 5m00 experimental structure highlighting side chains for the antigen residues and TCR residues receiving, on average, the highest tercile of attentions per chain (Figure 4D, Supplementary Figure 7). Residues with greater attention, particularly those in the TRB chain (which is more proximal to the antigen than the TRA), are frequently directly in contact with the antigen sequence.

### TCR-BERT can facilitate TCR design

Beyond achieving state-of-the-art results in classification and clustering, TCR-BERT enables novel computational approaches to important experimental and clinical challenges involving TCRs. Among these, one exciting domain is TCR engineering, which seeks to redirect T cell specificity by introducing synthetic TCR sequences into T cells. In principle, this strategy can boost T cells’ ability to recognize a given antigen, thus strengthening the immune system’s defense against specific pathogenic or malignant (cancerous) cells ^45^. This approach holds great promise in clinical treatments with targets ranging from cancers ^46 47^ to chronic viral infections like hepatitis B ^12^ and human immunodeficiency virus type 1 ^48^. However, the process of designing synthetic TCR sequences remains challenging. We use TCR-BERT to develop a proof-of-concept computational approach to rapidly screening and refining such TCR sequences. Intuitively, we use TCR-BERT to drive an iterative *in silico* directed evolution process. We start with a set of TCR sequences with no observed binding to an antigen of interest and use a classifier (built using TCR-BERT) to rank these according to their predicted likelihood of binding. We use the sequences with the highest predicted binding likelihood to generate additional, similar sequences. This is done by sampling from TCR-BERT’s masked amino acid predictions for randomly hidden positions in each TCR, thus leveraging TCR-BERT’s learned language model to introduce TCR residue mutations traversing the complex manifold of endogenous TCRs. We repeat the ranking and sampling steps using these newly generated sequences as input, iterating until we arrive at a set of sequences with desirable predicted binding (Figure 5A, see Methods for additional details). Importantly, this process does not simply identify the strongest binders within its input, instead using that information to generate novel TCR sequences.

**Figure 5:**
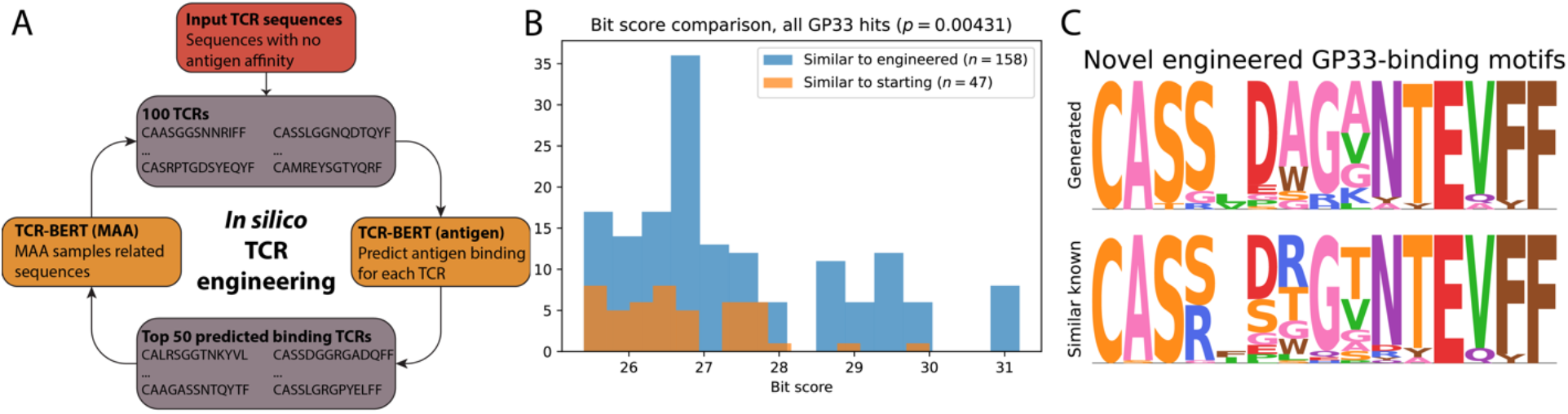
TCR-BERT enables in-silico engineering of novel TCR sequences. (A) To engineer sequences with affinity for the GP33 antigen, we take 100 endogenous TRA and TRB pairs with no measured binding to GP33, use a classifier based on TCR-BERT to select the sequences most likely to bind, and use those sequences to sample new TCR sequences using TCR-BERT’s masked amino acid predictions as a generative model. This directed evolution process is repeated to iteratively refine the pool of TCRs. This enriches for desirable binding properties and goes beyond simply choosing from the original inputs (Supplementary Figure 8). (B) To evaluate the efficacy of our TCR engineering process, we use BLAST to match both our starting and final engineered TRB sequences against previously identified murine TRBs. We find that final engineered sequences produce more significant hits to prior known GP33-binding TRBs (blue) compared to the starting set of TRBs (which also contains fewer hits). This is indicated by their significantly elevated bit scores, which indicate the log-size of the database needed to produce a comparable hit by random chance (displayed p-value, Mann-Whitney test). (C) Our *in silico* engineered sequences match several known GP33 binders that were not similarly matched by our starting sequences. These novel matches indicate that our TCR engineering process generates new yet biologically meaningful sequences. Here, we summarize these previously unobserved matches (bottom motif) as well as our generated sequences that bear significant similarity (top motif). These matches are individually illustrated in Supplementary Figure 9. We further show that we can generate additional motifs using different starting conditions (Supplementary Figure 10), thus demonstrating the flexibility of our approach.

We apply this procedure to engineer TRA-TRB pairs targeting GP33 binding using the version of TCR-BERT fine-tuned to predict LCMV GP33 binding from TRA-TRB pairs. We start with 100 TRA-TRB pairs with no measured binding drawn from the test set to ensure that TCR-BERT has never seen these sequences. We apply the directed evolution process described above and observe that the predicted GP33 affinity grows with successive iterations, reaching a minimum 95% predicted probability of binding after 7 iterations (Supplementary Figure 8A). These 50 generated TCRs share no clear sequence similarity to training examples with observed GP33 binding (Supplementary Figures 8B, 8C), indicating that our procedure generates novel TCRs not seen by the model.

To show that we are generating reasonable candidates for binding GP33, we use BLAST ^49^ to match both our input TRB set and final engineered TRB set against all known murine TRB sequences (we could not find adequate data to similar evaluate TRAs, see Methods for additional details). TRBs in the final set not only produce more hits to TRBs previously found to bind GP33 in an experiment never shown to TCR-BERT, but these hits are also more specific (indicated by significantly higher bit scores, Mann-Whitney test, *p* = 4.31 × 10^−3^, Figure 5B). Additionally, our engineered TRBs produce matches to 11 known GP33 binders that were not matched by our starting set (Figure 5C, Supplementary Figure 9); these further illustrate that our TCR engineering process generates novel sequences.

We additionally ran our TCR engineering procedure using a different set of starting sequences similarly drawn from non-binding test examples to further explore our procedure’s flexibility and generalizability (Supplementary Figure 10A). As before, we use BLAST to match the starting and final engineered TRBs against known murine TRBs and observe that our engineered sequences contain proportionally more GP33-binding sequences (Fisher’s exact test, *p* = 1.17 × 10^−4^). The engineered sequences are distinct from previous sequences (Supplementary Figure 10B), indicating that TCR-BERT is capable of flexibly generating a range of motifs that consistently mirror known biological motifs with GP33 affinity. While this specific usage of TCR-BERT as a platform for TCR engineering requires extensive follow up work and experimental validation, we showcase it to demonstrate how TCR-BERT can be leveraged as a computational platform for future technologies.

## Discussion

TCR-BERT is a large language model trained to model and embed T-cell receptor sequences. Compared to prior works using machine learning to predict TCR behavior, TCR-BERT is uniquely designed to leverage unlabeled data to learn a more general representation of TCRs, before being applied to or fine-tuned on specific downstream tasks. By leveraging this approach of unsupervised transfer learning, TCR-BERT enables state-of-the-art prediction of TCR-antigen binding and consistent, reliable grouping of TCRs likely to share antigen specificities. We further demonstrate that TCR-BERT enables these broad advances by focusing on structurally relevant residues with known biological importance for binding, despite being trained on only raw sequences that do not intrinsically carry this information. This suggests that TCR-BERT is likely learning general patterns across TCRs. Finally, we show how TCR-BERT can enable novel applications that go beyond canonical tasks like classification and clustering, such as computationally engineering TCR sequences with desirable binding characteristics.

There are several directions for improving and extending TCR-BERT. Most directly, our proof-of-concept TCR engineering work requires extensive follow-up experimental validations. More broadly, TCR-BERT does not leverage VDJ gene usage information in its design. Previous works have found that these additional annotations can improve antigen binding prediction performance compared to using TCR sequences alone ^29^, but the optimal approach for integrating these annotations into a large language model is an open question. From a usability standpoint, it can be difficult to understand exactly how TCR-BERT embeds or classifies a TCR sequence, especially when compared to traditional methods built around more transparent metrics like sequence similarity. Future work in model interpretability could alleviate this. Architecturally, research studying transformers is still rapidly evolving, so new methods for designing and training these large language models may be applicable to refining TCR-BERT. Researchers have also recently developed methods to use transformer models like TCR-BERT to study the evolution of protein sequences ^50^ – applying these to TCR-BERT may yield novel biological insights into the evolution and selection of TCRs ^51^. Finally, TCR-BERT would also greatly benefit from additional training data, especially as new technologies emerge that improve scalability of profiling TCR sequences along with their antigen specificities ^52^. Such data could also be used to extend TCR-BERT to include not just alpha/beta dimers, but gamma/delta dimers as well.

Beyond its direct contributions to antigen binding prediction and TCR clustering, we believe TCR-BERT represents a conceptual advance in the computational analysis of TCR sequences. Rather than training a new model from scratch for each specific antigen, TCR-BERT is a single model that can serve as a robust starting point for a wide variety of downstream tasks. This simultaneously reduces the amount of data required to build effective models, while lessening the amount of time and energy spent designing customized features, resulting in improved models that are easier to develop and share. It is our hope that this leads to more exploration of innovative applications for our growing computational understanding of TCR sequences, particularly in how we can leverage this understanding to inform our understanding of T cell function, and how we can ultimately apply these developments to enhance immunotherapies and patient care.

## Materials & Methods

### Datasets & preprocessing

From the pan immune repertoire database (PIRD) ^37^, we use the TCR-AB database containing 47,040 TRB sequences and 4,607 TRA sequences. Among these, 605 examples are explicitly paired TRA and TRB, and 8,429 examples have annotated antigen specificities. These annotated antigen specificities span 73 unique antigens. We use the PIRD dataset for masked amino acid modelling pre-training, and its labels for antigen classification pre-training and antigen cross-validation.

VDJdb is a curated dataset of T-cell receptor sequences with known antigen specificities ^36^. This dataset consists of 58,795 human TCRs and 3,353 mouse TCRs. More than half of the examples are TRBs (n= 36,462) with the remainder being TRAs (n= 25,686). Although these sequences are all labelled with a known antigen binding partner, we only used this dataset during the masked amino acid pre-training step. We empirically found that including this dataset’s labels did not improve the usefulness of TCR-BERT’s embedding for downstream classification or embedding tasks (data not shown).

The TCRdb dataset ^40^ consists of 139,00,913 TRB sequences of unknown antigen binding affinity. While we attempted to leverage TCRdb in our MAA pre-training step, we empirically found that doing so greatly increased training runtime without yielding improved downstream results (data not shown). Rather, we use TCRdb as a pool of unseen, naturally occurring human sequences of unknown antigen binding affinity. We sample from these sequences to create a “negative” set of TCR sequences. This is useful for building classifiers from datasets that only describe TCR sequences with known binding affinity (i.e., a positive label), but does not describe TCR sequences with no known binding affinity (i.e., a negative label). These negative sets are sampled at a ratio of 5 TCRdb negatives to each known positive example. This ratio is a round value that approximates the proportion of positive examples in the relatively exhaustive LCMV dataset, discussed below.

The murine LCMV GP33 dataset consists of T cells from the lung, liver, and spleens of mice infected with either LCMV Armstrong or LCMV Clone 13. Following tissue dissociation, single cell suspensions were stained with class I tetramer H-2Db LCMV GP33-41 (KAVYNFATC) (PE) and then sorted via flow cytometry as tetramer high, mid, or negative. TCR sequencing was performed using the 10X 5’ Single Cell Immune Profiling Solution Kit (v1.1 Chemistry) using the 10x Chromium Single Cell V(D)J Enrichment Kit for mouse T cells, according to the manufacturer’s instructions. Single-cell TCR-seq libraries were sequenced on an Illumina NovaSeq S4 sequencer using the following read configuration 26bp Read1, 8bp i7 Index, 91bp Read2. TCR reads were aligned to the mm10 reference genome and consensus TCR annotation was performed using cellranger vdj (10x Genomics, version 3.1.0). We keep only unique TRA/TRB clones where 80% of annotated cells assigned the same label, in which the clone is assigned the majority label. Overall, this results in (n=17,702) unique TRA/TRB pairs with consistent labels that we use for model training and evaluation. Among these, 13% (n=2306) are observed to have mid or high binding – we consider these “positive” examples of TRA/TRBs binding GP33. This dataset is also being used for a separate, unrelated manuscript under preparation for submission; this manuscript contains more details surrounding exact experimental conditions for gathering this data as well as details on data access.

For all TCR sequence records, we exclude any examples that include residues outside the set of 20 standard amino acids, e.g., wildcard residues that indicate variability.

### Modelling, pre-training, and layer selection

TCR-BERT is implemented in Python primarily using the PyTorch ^53^ and Transformers libraries ^54^. TCR-BERT uses a lightly modified version of the BERT language modelling architecture. We provide a brief description of BERT and general transformer models here; for full details, please refer to the original BERT manuscript ^31^. Transformer models ^55^ use a series of blocks that apply attention and feed-forward layers. Attention attempts to model pairwise interactions in an input sequence by learning how strongly the embedding representation of each token should be influenced by the embedding of other tokens (including itself). Applying several layers of such blocks allows models like BERT to learn increasingly complex interactions between tokens that ultimately capture higher-order concepts like grammatical structure. These transformer networks have been shown to outperform more conventional recurrent and convolutional models in various natural language problem settings.

TCR-BERT’s input is TCR sequence of length *M*, formatted as a series of *M* tokens spanning the set of 20 amino acids. These input tokens are then padded with special tokens: a classification token *C* as a prefix, and a separator token *S* as a suffix. The padded input tokens are then passed through a trained embedding layer that maps each token (amino acid) to a continuous representation of 768 dimensions. As this token embedding does not capture positional information, our TCR-BERT model follows the BERT model in adding a positional encoding to the amino acid embedding. This summed sequence embedding is then fed through a series of 12 transformer blocks to arrive at the overall sequence embedding. This sequence embedding represents each input amino acid as a (*M* + 2) × 768 matrix. The sequence embedding can then be fed into various “heads” that perform pretraining and downstream tasks such as masked amino acid prediction or sequence classification.

We pre-train TCR-BERT using two objectives optimized sequentially. First, we pretrain using a masked amino acid (MAA) modelling objective, where we randomly hide, or “mask” 15% of the amino acids in each TCR amino acid sequence in the training set, and train TCR-BERT to predict these masked amino acids. Architecturally, this is done by appending a MAA “head” network to the previously described transformer network. This MAA head is a simple, fully connected layer that maps TCR-BERT’s per-residue hidden representation to logits (20 dimensions, corresponding to each amino acid), followed by a softmax activation. MAA pretraining is done using both TRA and TRB sequences from the VDJdb and PIRD datasets and aims to leverage the large amount of TCR sequences with (potentially) unknown antigen specificity to learn the “grammar” of a valid TCR sequence. TRA and TRB sequences are given to the model without features or flags distinguishing the two. We use a random 85/15 train/test split for MAA pre-training and tune hyperparameters based on test set loss. We perform grid search across the following hyperparameters with final chosen values in **bold**. Several of these hyperparameters describe architectural configurations (e.g., dimension of the hidden representation) whose values apply to downstream training/tasks as well. Default values for the BERT architecture are indicated. Hyperparameters and architectural configurations not indicated are left at their default values.

- Hidden representation dimensionality: [144, 384, **768**] (BERT default: 768)
- Intermediate representation dimensionality: [**1536**, 3072] (BERT default: 3072)
- Number of attention heads: [6, 8, **12**] (BERT default: 12)
- Number of transformer layers: [8, **12**] (BERT default: 12)
- Batch size: [128, **256**]
- Learning rate warmup (number of training steps to linearly “ramp up” learning rate to specified value): [**0**, 0.1]
- Training epochs: [10, 15, 25, **50**, 100]
- Learning rate: [2e-5, **5e-5**]

We also reduce the maximum positional embedding length to be 64 (as opposed to BERT’s default maximum of 512) to reflect the relatively short TCR sequence lengths compared to sentence lengths in natural language. We use linear learning rate decay to 0 over training epochs, coupled with the AdamW ^56^ optimizer and negative log likelihood loss.

The second objective is a multi-class classification task where we train TCR-BERT to classify each input TRB sequence as binding to a one of a set of profiled known antigens. We focus on TRB sequences exclusively, as very few TRA sequences have annotated antigen specificities (and even fewer paired TRA-TRB examples have such annotations). To train this objective, we subset the antigens represented in the PIRD dataset to include only antigen sequences with at least 6 positive examples; antigens with fewer positive examples are aggregated into a single “other” label. This results in 44 antigen labels and 1 “other” antigen label for a total of 45 labels, distributed among with 6,235 TRB sequences. These are randomly split into training, validation, and test sets using a 70/15/15 split. The antigen classification “head” consists of a “pooling” layer (a feedforward network projecting the 768-dimensional classification token embedding into 768 dimensions with Tanh activation) and a “classifier” layer (a feedforward network projecting the 768-dimensional output of the pooling layer into the number of labels). As this pre-training objective predicts a single antigen for each TRB, we use a softmax activation coupled with a negative log likelihood loss. For this pre-training objective, we perform grid search over the following hyperparameters optimizing for validation set AUPRC, with selected values in bold:

- Learning rate: [2e-5, **5e-5**]
- Batch size: [**128**, 256]
- Learning rate warmup: [0, **0.1**]
- Training epochs: [10, 15, **25**, 50]

We additionally perform early stopping after no improvement in validation set AUPRC after 5 epochs.

After these pre-training steps, we select a representation layer from TCR-BERT that is most conducive to downstream tasks like clustering or building classifiers. This is necessary as empirical works have found that the last layer of large language models like BERT are often too specialized for pre-training tasks to be generally useful for embedding new inputs. To do this, we sweep across the last seven layers of the model, use each layer to train a PCA-SVM classifier (see following section for full details) using the LCMV GP33 dataset (using a random 70/15/15 train/valid/test split), and select the layer maximizing validation set AUPRC. We repeat this applying various levels of subsampling to the training data to choose a layer that consistently performs well regardless of the amount of data available. We find that the 8^th^ layer is optimal, and we use this layer for all experiments that use TCR-BERT to generate embeddings (e.g., PCA-SVM, Leiden clustering, etc.), including for datasets other than LCMV.

### Downstream classifiers

After pre-training, we leverage TCR-BERT’s learned representation using various downstream classifiers. One approach involves building classifiers directly on the embedding that TCR-BERT produces, treating TCR-BERT as a black box for generating TCR embeddings. We first average TCR-BERT’s representation of each amino acid in a TCR sequence (excluding special tokens like classification and separator tokens), using the optimal layer discussed above. We then use PCA to reduce the resulting representation’s dimensionality to the top 50 principal components (PCs) and train a support vector machine classifier with radial basis function kernel on this reduced embedding. This can theoretically be applied to either TRA or TRB sequence alone or both, though we focus on embedding the TRB sequence in our work. Using PCA to summarize the embeddings helps the SVM to better focus on major sources of variation in the embedding.

TCR-BERT can also be fine-tuned to perform classification. Rather than treating TCR-BERT as a fixed black box for generating continuous embeddings from discrete amino acid sequences, fine-tuning modifies TCR-BERT using the pre-trained parameters as a starting point. In our work, we fine-tune TCR-BERT to perform antigen binding prediction given a TRA and TRB sequence pair. Architecturally, this paired prediction model is comprised of two separate TCR-BERT transformers individually responsible for embedding the TRA or TRB sequence, respectively. Both are initialized using weights from the masked amino acid pretraining step, as MAA pre-training is agnostic of TRA/TRB identity. We extract the initial “classification” token embedding from both the TRA and TRB, concatenate the two embeddings, and apply a fully connected layer mapping to two outputs with softmax activation to generate binding predictions. Notably, training this overall model tunes each encoder towards patterns specific to either the TRA or TRB. Hyperparameters for this model are selected maximizing validation set AUPRC using grid search over the following values (final values are in bold):

- Learning rate: [5e-5, **3e-5**, 2e-5]
- Training epochs: [10, **25**, 50, 100]
- Dropout (in final fully connected layer, does not affect TCR-BERT itself): [0.1, **0.2**]

We train using a batch size of 128, linear learning rate decay over training epochs, and the AdamW optimizer. While we focus on fine-tuning targeting both TRA and TRB pairs, TCR-BERT could also be fine-tuned to target only TRB sequences using a similar approach without concatenating multiple embeddings.

In several of the experiments described in our work, we aim to build a classifier distinguishing human antigen binding TRB sequences from a random background, sampled at 5 background sequences per binding sequence. To create this random background, we randomly select TRB sequences from the TCRdb dataset ^40^, which contains human TRB sequences of undetermined antigen affinity (see above). This process attempts to mimic a natural sample of TCRs, where there are many “background” TCRs and a subset of TCRs with binding to a specific antigen. This ratio is similar to what is observed in tissues from severely diseased mice, though this ratio may be lower in humans.

To evaluate classifier performance, we primarily focus on area under the precision recall curve (AUPRC). AUPRC is a much more informative metric in cases of extreme class imbalance towards negative cases (i.e., when there are very few positive labels). Such class imbalance is commonly observed in the context of TCR specificity, where most TCRs will not bind to an antigen. We also present area under the receiver operating characteristic (AUROC) in some instances for completeness. The expected AUPRC for a random classifier is the proportion of positive examples the dataset contains, and the expected AUROC is 0.5.

### Downstream clustering

Our method for using TCR-BERT to embed TCR sequences for clustering analysis shares parallels with our method for building a SVM on top of TCR-BERT’s embeddings. As before, we use the previously identified optimal transformer layer and reduce the dimensionality of the embedding using PCA. We visualize the resulting embedding using Uniform Manifold Approximation and Projection (UMAP) ^38^ and cluster sequences using the Leiden clustering algorithm ^42^. To control the number and granularity of output clusters, we vary the resolution parameter for Leiden.

To evaluate clustering performance, we use two metrics originally developed to quantify the performance of the GLIPH algorithm ^15^: percent clustered and percent correctly clustered. Each group of TCR sequences with 3 or more TCR sequences is considered “clustered.” The number of such clustered TCRs divided by the total number of TCRs gives the percent clustered. To calculate percent correctly clustered, we iterate over each cluster and for each TCR in that cluster, we retrieve its associated label (i.e., the antigen that that TCR binds to). If there is a dominant (≥50% occurrence) label within the cluster, the entire cluster is assigned that dominant label. Percentage correctly clustered represents the average accuracy of each clustered TCR evaluated against this majority label. As a toy example, consider a single cluster with sequences [*a, b, c, d, e*] and corresponding antigens labels [*x, x, x, x, y*]. The percent correctly clustered is 80%, as this would be the accuracy if we had assigned the *x* label to all points in the cluster.

To evaluate clustering runtime, we use the UNIX “time” command with TCR-BERT and GLIPH methods. Timing includes the entire process from reading a new input of TRB sequences, calculating clusters, and writing relevant output files. Runtime benchmarking is done with different subsets of murine LCMV GP33 TRBs. All benchmarks are run using the same machine with an Intel i9-9960X processor, 128GB of RAM, and an Nvidia GeForce RTX 2080Ti GPU. No other foreground processes were run during benchmarking.

### Evaluated external methods

We use various tools to contextualize TCR-BERT’s ability to perform classification and clustering. GLIPH was downloaded from the authors’ GitHub repository: https://github.com/immunoengineer/gliph. To cluster our TRB sequences, we run GLIPH using the “group discovery” script using varying values for the global convergence cutoff. We attempted to use the updated GLIPH2 algorithm ^26^, but this tool requires additional inputs, including Vb and Jb usage, which makes direct comparisons with TCR-BERT difficult.

To evaluate against the Evolutionary Scale Model (ESM), we downloaded the ESM-1b model from the PyTorch model hub. To embed TCR sequences, we use the default mode of extracting the final transformer layer’s embedding and average the embedding for each amino acid to obtain the embedding for the overall sequence. This follows the recommendations given in the original authors’ GitHub repository https://github.com/facebookresearch/esm. These embeddings are then used for PCA-SVM.

The TAPE model ^34^ was downloaded from the authors’ GitHub: https://github.com/songlab-cal/tape. We use the UNIREP version of their model to generate averaged embeddings using the default configuration. These embeddings are then used for PCA-SVM.

The SETE model ^30^ does not provide a simple code interface to train and evaluate on a dataset; we instead used the authors’ manuscript and reference code from their GitHub repository: https://github.com/wonanut/SETE to re-implement their algorithm.

The DeepTCR model ^29^ version 2.0.10 was installed via Python pip. To evaluate DeepTCR’s performance on our GP33 dataset, we first train DeepTCR on the bundled murine antigen data using TRA/TRB featurization (i.e., excluding VDJ annotations for comparability). We then freeze DeepTCR’s parameters and use the resulting model to embed our LCMV GP33 TRA/TRB sequence pairs. These embeddings are then used as input to PCA-SVM classification. We also evaluated other classifiers such as logistic regression on top of these embeddings, but PCA-SVM yielded the best performance.

We developed an in-house convolutional architecture (ConvNet) as an additional baseline for predicting binary antigen binding given TCR sequences. This is motivated by the fact that many researchers have broadly demonstrated strong results using convolutional networks to perform classification and motif discovery within biological sequences ^29 57 58^. ConvNet maps amino acids to a 16-dimensional embedding, followed by convolutional layers mapping to 32, 32, and 16 channels with kernel sizes of 5, 5, and 3 respectively. The output of the final convolutional layer is then passed through a fully connected layer to a 2-dimensional output with softmax activation to predict antigen binding probability. ConvNet is trained using the Adam optimizer ^59^ with cross entropy loss, a learning rate of 0.001, and a batch size of 512 along with early stopping after 25 epochs of no improvement to validation AUPRC. When training on paired TRA/TRB sequences, we modify ConvNet to learn a separate embedding and convolutional portion for each chain. The final convolution embeddings are then concatenated and input to a single fully connected layer with softmax activation for output probabilities.

### Interpreting and contextualizing model attentions

To obtain TCR-BERT’s attention across TRA and TRB chains, we use the version of TCR-BERT fine-tuned to perform LCMV GP33 antigen prediction. Recall that this model contains two fine-tuned variants of the TCR-BERT model embedding TRA and TRB sequences, respectively. Thus, for each chain within the TCR, we examine the respective fine-tuned TCR-BERT variant and extract the model attentions from the first classifier token at the last transformer layer for each of the 12 attention heads. We then trim these attentions to the length of the input *L* (excluding padding tokens). This results in a matrix of shape 12 × *L* for each chain in each example. This approach is heavily inspired by the bertviz library ^60^.

We average these attention matrices across test set examples to paint a clearer picture of typical TCR-BERT attentions. For this, we restrict to test examples with the same length TRA and TRB (n=157, 12 and 14 residues for TRA and TRB, respectively). Test set examples are used as they are not seen for training or hyperparameter tuning, and thus represent a more robust evaluation that more closely describes potential real-world use. When contextualizing TCR-BERT’s attentions, we average across the 12 attention heads to obtain a vector of length *L*.

We contextualize TCR-BERT’s per-residue attentions using the antigen distance of each residue, computed from the 3D structure of the MHC-antigen-TCR complex. We define antigen distance as the minimum Euclidean distance between a given TCR residue and any residue in the antigen peptide. Each residue’s 3D coordinates are summarized as an average of the atoms comprising the residue to alleviate computationally intensive pairwise atom calculations. We apply this to Protein Data Bank (PDB, https://pdb101.rcsb.org/) ^61^ structures 5m00, 5m01, and 5m02. These three structures exhibit minor differences from our dataset. They study a slightly modified LCMV GP33 antigen (KAVANFATM versus our antigen sequence KAVYNFATC) interacting with the TRA/TRB pair CAALYGNEKITF/CASSDAGGRNTLYF (also of lengths 12 and 14 residues, respectively). This specific TRA/TRB pair is not predicted to bind to our GP33 antigen, but it is unclear whether this is driven by the differences in the specific antigen or by model error. We believe that our results should be robust despite these minor differences, as the overall structure of these interactions should be similar.

### TCR engineering

Our *in silico* TCR engineering process begins with a set of starting sequences with no binding affinity for the antigen in consideration. In our case, this consists of 100 non-binding TRA/TRB pairs randomly selected from the LCMV GP33 test set. Selecting from the test set ensures that the model has not been trained or tuned on these specific sequences, as would be the case for real-world usage. We then give these TRA/TRB pairs to the version of TCR-BERT fine-tuned to predict LCMV GP33 binding form TRA/TRB pairs, ranking them by their predicted GP33 binding. We take the top half of these sequences (n=50) and use them as seeds to generate a set of new sequences (n=100).

To sample a new TRA/TRB pair, we start by randomly selecting one of the given seed sequence pairs. We then use TCR-BERT to mutate both the TRA and TRB, introducing two amino acid mutations to each chain. Mutations are introduced incrementally by choosing a single random position within the chain, masking that position, and giving the masked input to TCR-BERT (pretrained on MAA only) to predict the masked amino acid. This yields a probability distribution describing the most likely amino acids given the rest of the sequence. We rely on this prediction to explore the “grammar” of valid TCRs, such that we do not generate sequences that are untenable (e.g., a sequence entirely consisting of a single repeated residue). We randomly sample from the top 5 amino acids in this distribution.

These (n=100) generated TCR chains are then given to TCR-BERT to re-rank, and the process of using the top sequences to re-generate new TCR chains with (hopefully further) enhanced binding is repeated. This cycle is iterated until the predicted binding converges to a satisfactorily high value.

We use BLAST ^49^ to check the resulting TRB sequences against known murine TRB sequences. We construct a custom BLAST database using all protein sequences from RefSeq protein matching the query string “t cell receptor beta chain[All Fields] AND “Mus musculus”[porgn]” (n=2467). We then use BLAST version 2.5.0 to match sequences against this database using an E-value cutoff of 0.001. For a baseline comparison, we ran the top 50 predicted GP33 binding starting sequences through BLAST as well against this same database. We compared the number of resulting GP33-related hits versus other, non-related hits for the starting and final BLAST matches using Fisher’s exact test. We additionally compare the GP33-related bit values (proportional to the log-size of the database required to produce such a hit by chance) corresponding to matches to the starting set, and matches to the engineered set, using a Mann-Whitney test.

### Miscellaneous external libraries

Baseline neural networks and fine-tuned TCR-BERT models were developed using version 0.10.0 of the skorch library. 3D protein structures are visualized using PyMOL (The PyMOL Molecular Graphics System, Version 2.5 Schrödinger, LLC.). Motif logos are generated by using MUSCLE (version 3.8.1551) ^62^ to generate a multiple sequence alignment that is visualized using the Logomaker Python package ^63^. All other plots were generated using the matplotlib ^64^ and seaborn libraries. All metrics were computed using the scipy ^65^, scikit-learn ^66^, and numpy ^67^ libraries. We additionally use scanpy (version 1.7.1) and anndata (version 0.7.5) to simplify implementation of select clustering analyses ^68^.

### Code and model availability

Code implementing the TCR-BERT model and all analysis (antigen classification, clustering, TCR engineering, and model attention interpretation) is available from GitHub at https://github.com/wukevin/tcr-bert. Trained models are publicly available from the HuggingFace online model hub. Specifically, the version of TCR-BERT pre-trained on MAA and antigen classification is available at https://huggingface.co/wukevin/tcr-bert, and the version of TCR-BERT pre-trained on MAA only is available at https://huggingface.co/wukevin/tcr-bert-mlm-only. Please refer to our GitHub repository and the HuggingFace transformers library (https://huggingface.co/transformers/index.html) for complete usage details.

## Author contributions

H.Y.C., J.Z., K.E.W., and K.E.Y. defined the original motivations for this research. K.E.W. designed and performed all modelling, model benchmarks, and model applications. K.E.Y., B.D., J.A.B., Y.X., T.E., and A.S. performed data collection and preprocessing as well as access to novel data. All authors provided input for interpreting results and contributed to writing the paper.

## Acknowledgement

H.Y.C. is supported by RM1-HG007735 and Parker Institute for Cancer Immunotherapy. H.Y.C. is an Investigator of the Howard Hughes Medical Institute. T.E. is supported by R01-AI130152, R21-AI161040, and the Leukemia and Lymphoma Society Scholar Award. J.A.B was supported by a Stanford Graduate Fellowship and the National Science Foundation Graduate Research Fellowship under Grant No. DGE-1656518. K.E.Y. was supported by the National Science Foundation Graduate Research Fellowship Program (NSF DGE-1656518), a Stanford Graduate Fellowship, and an NCI Predoctoral to Postdoctoral Fellow Transition Award (NIH F99CA253729).

## Supplementary Figures

**Supplementary Figure 1:**
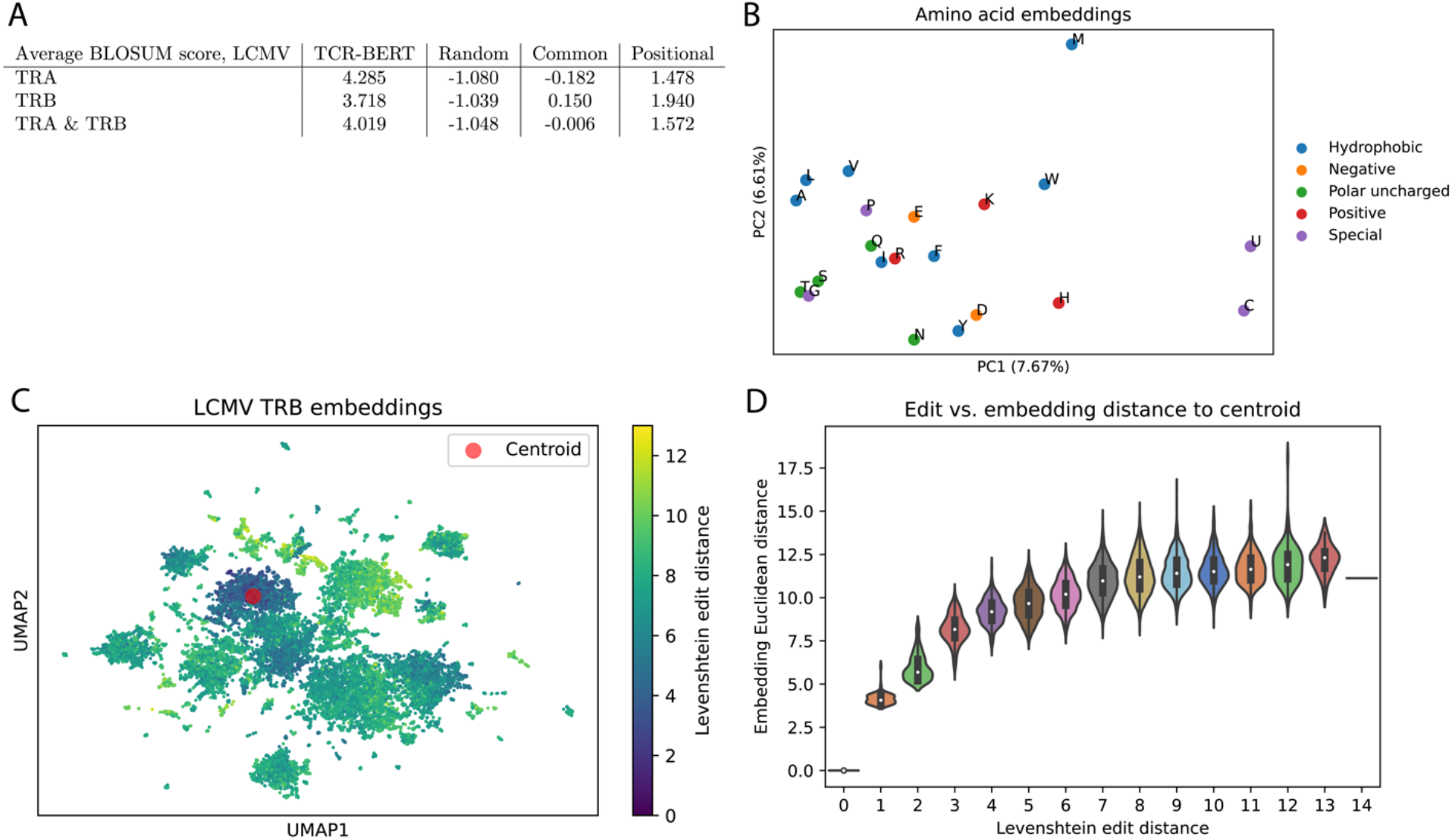
Validating TCR-BERT pretraining. (A) TCR-BERT’s performance on masked amino acid prediction on an in-house LCMV GP33 murine dataset, which is not used for pre-training. One amino acid is masked from each input sequence, and the TCR-BERT’s top prediction is scored against the hidden amino acid using the BLOSUM62 ^69^ scoring matrix. Higher values indicate more biologically similar predictions. We compare TCR-BERT’s performance to three baselines: predicting a random amino acid (random), predicting the most common amino acid regardless of positional information (common), and predicting the most common amino acids at each position (positional). This positional baseline is conceptually similar to using a multiple sequence alignment to generate predictions. TCR-BERT outperforms all baselines for predicting masked amino acids, which suggests that it successfully captures complex patterns in the grammar of TCR sequences. (B) TCR-BERT’s learned embeddings for each of the 20 amino acids, visualized using PCA. Points are colored by biochemical properties and labelled using standard one-letter abbreviations. We observe separation according to biochemical properties of these amino acids, such as by hydrophobicity along the x-axis. Biochemically similar residues also appear to have similar embeddings, e.g., alanine (A), leucine (L), and valine (V). (C) UMAP visualization of TCR-BERT’s embedding of the TRB sequences in the LCMV dataset. Each point represents one TRB sequence and is colored by Levenshtein edit distance to the randomly chosen “centroid” sequence (red). TCR-BERT embeds similar sequences in similar space; further sequences are more dissimilar as reflected by larger edit distances. Since sequence similarity is a known heuristic for predicting TCR binding, this suggests that TCR-BERT’s embedding is conducive to downstream TCR sequence analyses. (D) Plot comparing Levenshtein edit distance between TRBs relative to the (red) centroid TRB shown in (C) and their Euclidean distance in TCR-BERT’s embedding space. X-axis denotes the (discrete) edit distance to the centroid, and y-axis denotes the distribution of (continuous) Euclidean distances for all TRBs with that edit distance. We observe a strong correlation between edit and embedding distance, which lends additional support to the observation made in (C).

**Supplementary Figure 2:**
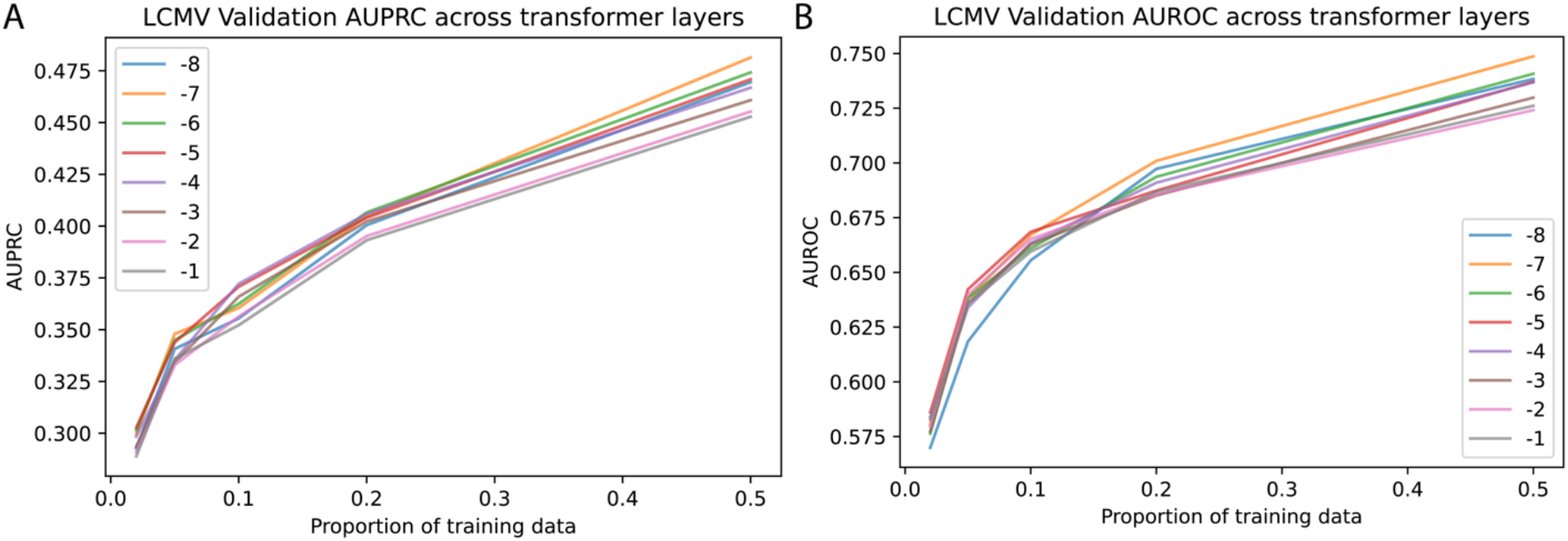
Layer selection for embedding. To choose the optimal representation within TCR-BERT for downstream tasks, we choose a layer that maximizes validation set performance within the LCMV GP33 dataset. We sweep across the last 8 transformer layers’ outputs, evaluating AUPRC (A) on the validation set (fixed size) for various levels of available training data from (x-axis). We find that the 8^th^ layer (−5 in above plots) most consistently produces the best AUPRC (our primary evaluation metric) across the widest range of training data sizes. We sanity check that this layer provides reasonable AUROC values (B) as well. We use this representation for all results, analyses, and visualizations that do not involve fine-tuning TCR-BERT. Note that we perform this layer selection once and do not repeat or adjust it for other datasets or other applications (e.g., TCR clustering).

**Supplementary Figure 3:**
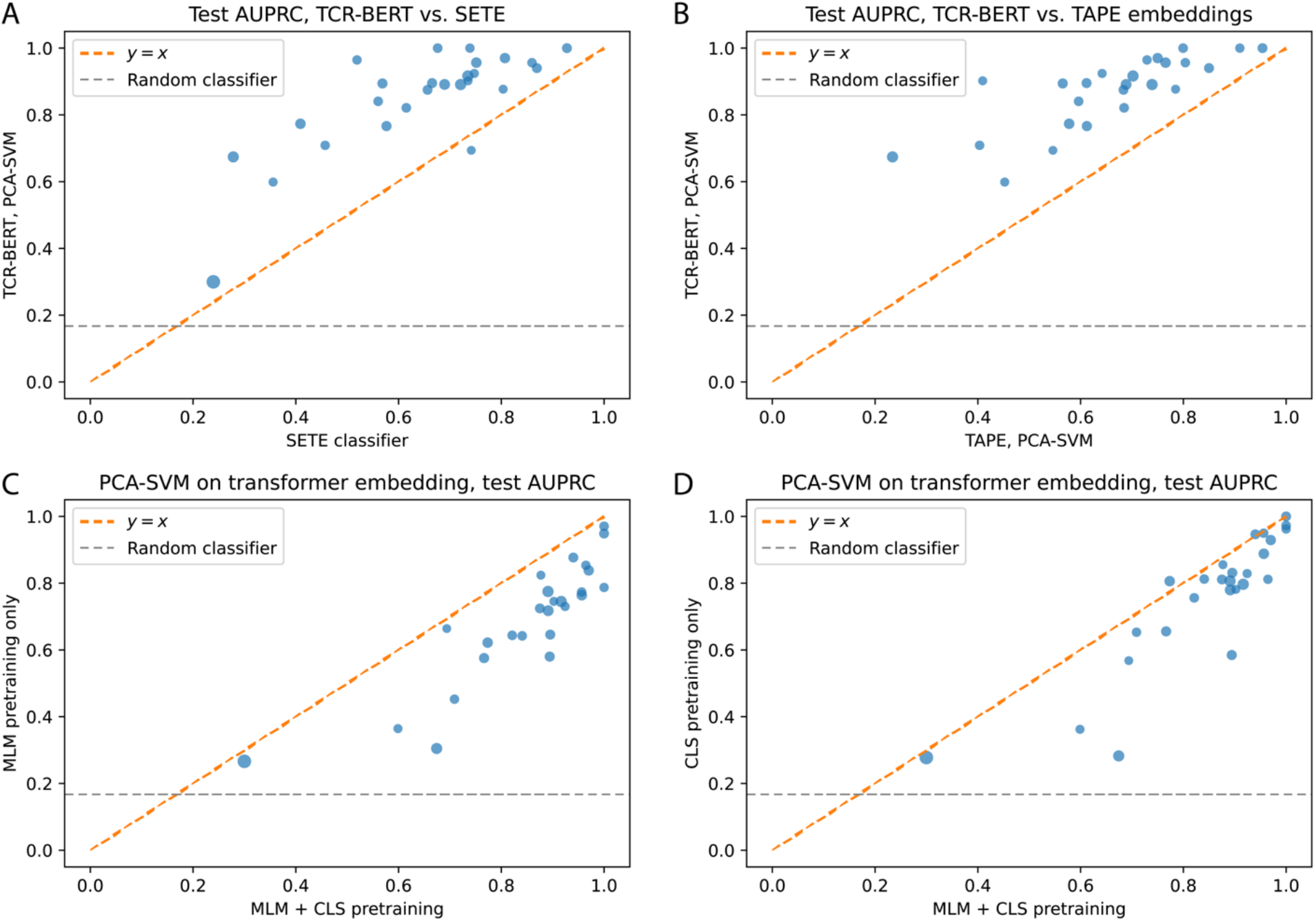
Additional antigen cross-validation performance comparisons. In each experiment, we take one of 26 antigens and its associated binding TRBs and spike in background human TRBs at a ratio of 5:1. We evaluate test AUPRC using a random 70/30 train/test split. In each panel, the grey line indicates performance of a random classifier, and the orange line indicates equal performance between the compared methods. Points above the orange line indicate instances where the y-axis model performs better, and vice versa. (A) Compares SETE, a supervised tree-based model (x-axis) to PCA-SVM on TCR-BERT’s embedding (y-axis). In 25 of 26 antigens, TCR-BERT exhibits improved test set AUPRC (p=4.67e-06, Wilcoxon test). (B) Compares performance of PCA-SVM using TAPE’s embeddings (x-axis) versus TCR-BERT’s embeddings (y-axis). In every tested case, TCR-BERT provides superior performance (p=7.32e-05, Wilcoxon test). This test isolates the effect of using different protein language models to embed TCRs while keeping the classifier module constant. Along with Figure 2B, this indicates that TCR-BERT outperforms general purpose protein language models for TCR embedding and modelling. (C) We additionally evaluate performance when omitting each of our two pre-training steps. Here, we leave out the classification pre-training step (i.e., using MAA only, y-axis), comparing it to the TCR-BERT model with both pretraining steps (x-axis). Omitting classification pre-training results in a drop in performance in all cases (p=4.15e-06, Wilcoxon test). (D) We similarly evaluate the full TCR-BERT pre-training (x-axis) against using only classification pre-training (i.e., no MAA, y-axis). We observe a decrease in performance in 24 of 26 cases (p=1.61e-05, Wilcoxon test). Along with panel (C), these results indicate that both of our pre-training steps are necessary for enabling TCR-BERT’s strong performance.

**Supplementary Figure 4:**
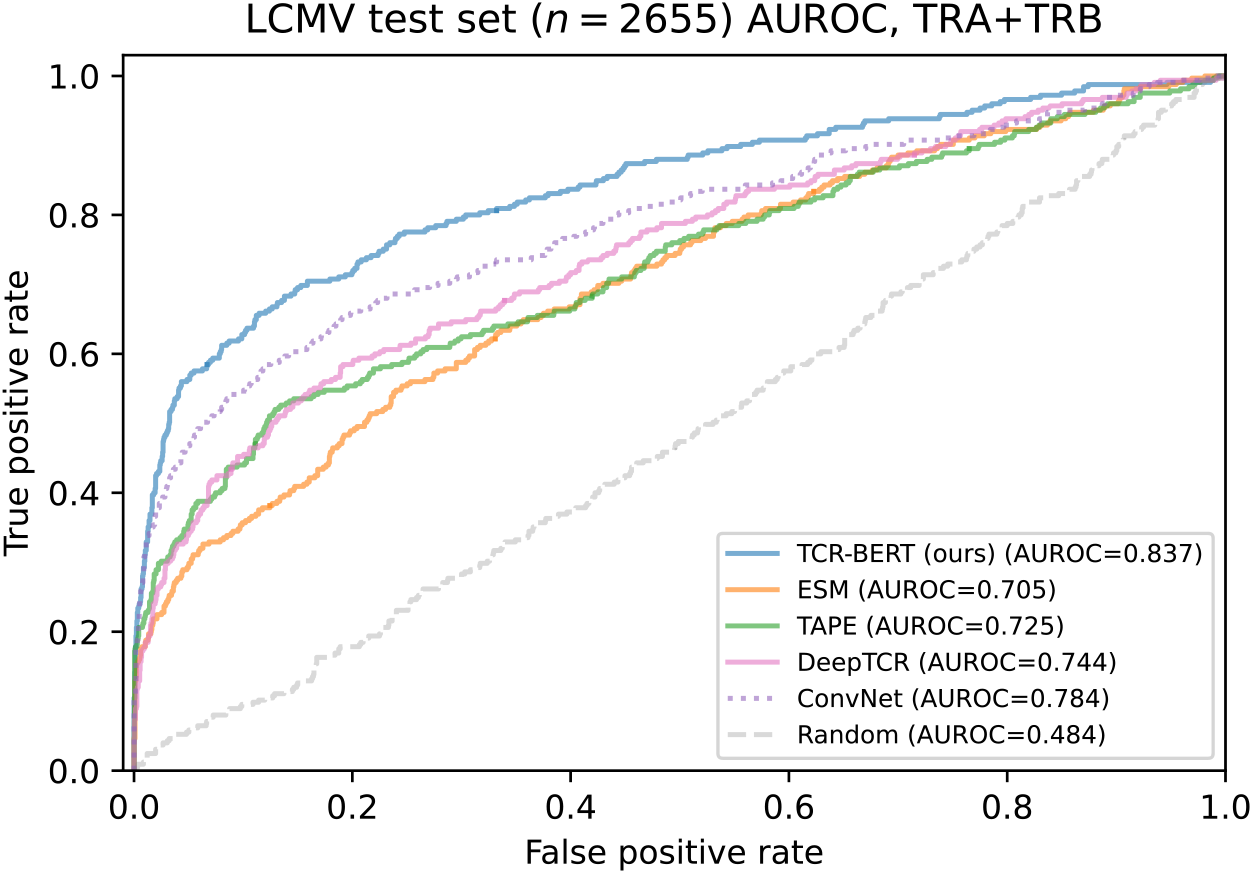
LCMV test set AUROC. AUROC curves describing various classifiers’ test performance predicting GP33 binding given paired TRA/TRB sequences. TCR-BERT provides the best performance when evaluated on AUROC (shown here) as well on AUPRC, our primary evaluation metric (Figure 2D). Solid line indicates models leveraging pre-training, whereas dashed lines indicate models that use only supervised training.

**Supplementary Figure 5:**
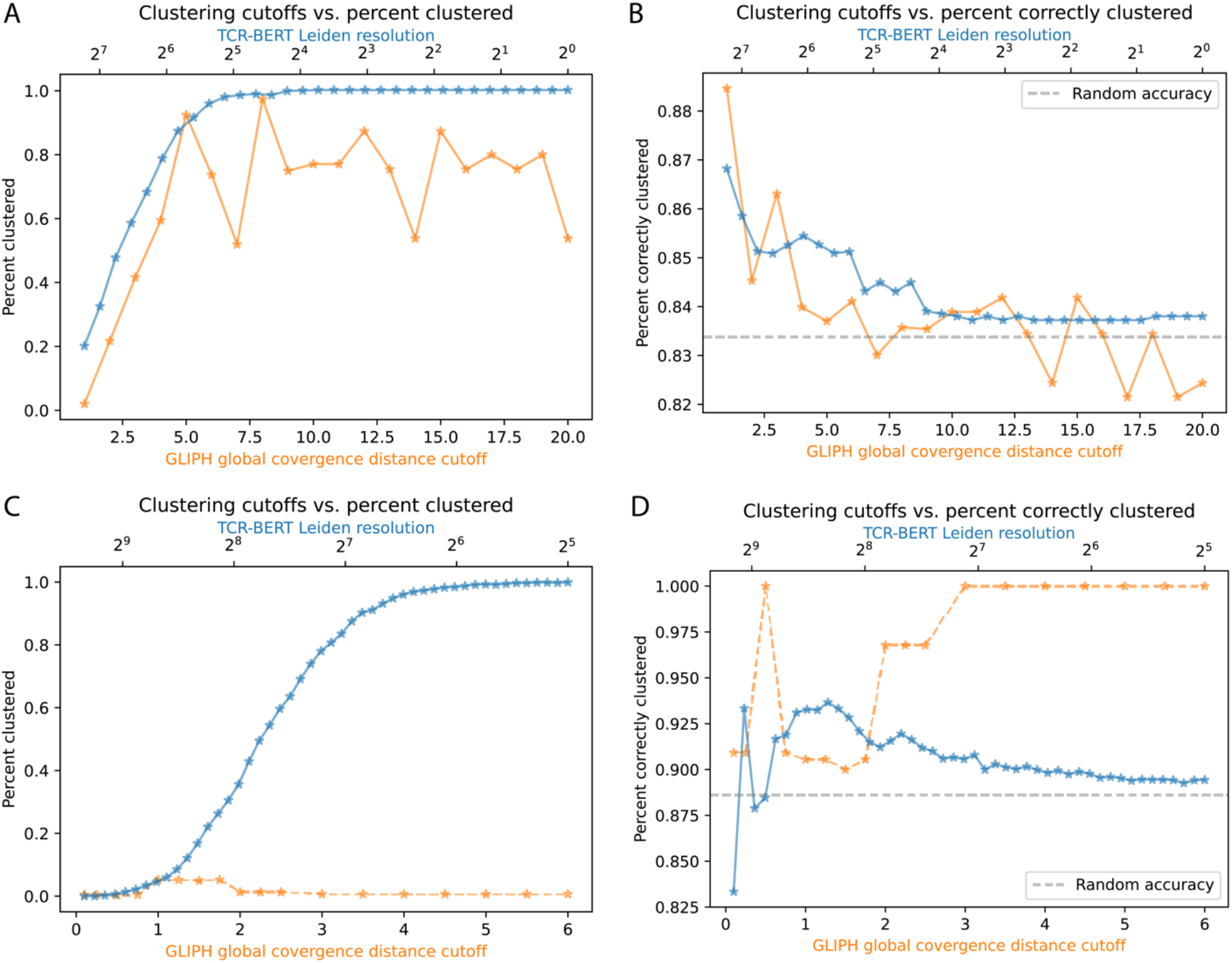
Detailed comparison of GLIPH and TCR-BERT antigen specificity groupings. Both GLIPH and TCR-BERT produce TRB groupings with configurable degrees of granularity. This is controlled using the resolution parameter to the Leiden clustering algorithm for TCR-BERT (larger values are more granular) and the global convergence distance cutoff for GLIPH (smaller values are more granular). We compare the performance of TCR-BERT and GLIPH across various clustering granularities using human NP177 antigen (A, B) and murine LCMV GP33 (C, D). (A) Shows percent clustered for a mixture of human NP177 antigen for TCR-BERT (blue) and GLIPH (orange). TCR-BERT converges to having all sequences clustered as the clustering becomes less stringent, while GLIPH produces variable, unpredictable results. (B) Shows the percent correctly clustered for these two methods, again for the NP177 antigen. Both methods trend towards lower percent correctly clustered as clustering granularity decreases. However, TCR-BERT’s clustering stabilizes above the expected correctness of a random classifier, whereas GLIPH’s behavior is much less consistent, frequently dipping below random. (C) Shows percent clustered for clustering methods on the murine LCMV GP33 dataset. With TCR-BERT, looser clustering configurations allow for clustering of the entire dataset (blue) as it does for the NP177 dataset (panel A). GLIPH on the other hand, produces no consistent relationship between its configuration and percentage clustered (orange). Indeed, GLIPH struggles to cluster any meaningful proportion of the given TRBs. This reveals that GLIPH not only exhibits erratic behavior across granularity configurations, but also behaves inconsistently when evaluated on different datasets. (D) Shows percent correctly clustered for these methods on the LCMV GP33 dataset. As before, TCR-BERT produces more predictable performance (blue). However, there are a few cases where TCR-BERT does poorer than random. These only occur when TCR-BERT clusters only a handful of sequences, when a single misclassified sequence has an outsized impact on correctness; as TCR-BERT clusters more sequences (lower resolution values, top horizontal axis), it consistently stays above random accuracy. While it might appear that GLIPH (orange) achieves perfect accuracy towards higher cutoffs, this is misleading as those cutoffs also lead to extremely few sequences being clustered (panel C), which vastly reduces the usefulness of the small handful of correctly clustered sequences. Increasing GLIPH’s global convergence distance cutoff seems to increase the percent correctly clustered in this case, which is contrary to the behavior observed for the NP177 human dataset (panel B). TCR-BERT’s behavior, on the other hand, is consistent across parameters for both these datasets. Overall, these results show that TCR-BERT exhibits consistent, predictable, and improved clustering behavior compared to GLIPH.

**Supplementary Figure 6:**
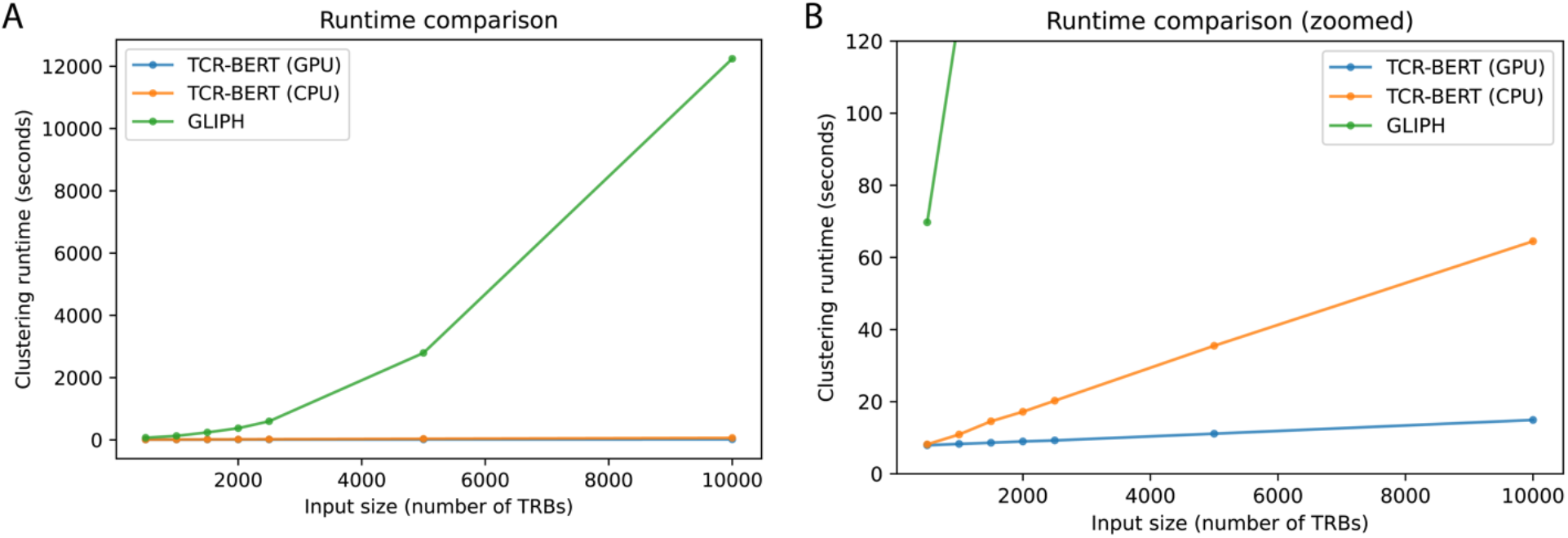
Runtime comparison for methods generating groupings for antigen specificity. Comparison of elapsed time, in seconds, required to cluster TRB sequences with predicted shared antigen specificity. All methods are benchmarked on the same LCMV GP33 murine TRB sequences, subsetted to various input sizes (x-axis), and are run on the same machine. TCR-BERT can be run with GPU acceleration or using only the CPU; both cases exhibit similar runtime performance. GLIPH does not support GPU acceleration. (A) Shows runtime performance as we scale from 500 to 10,000 input sequences. GLIPH’s runtime scales super-linearly. For example, increasing the number of input sequences by 4x from 2500 to 10000 results in a ∼21x increase in runtime from about 10 minutes to over 3 hours. This makes running GLIPH prohibitively time-consuming on larger datasets, especially when evaluating multiple parameters/configurations. (B) Truncates the y-axis to lower values to clearly show TCR-BERT’s runtime characteristics. Even without GPU acceleration, TCR-BERT can process 10,000 inputs in less time than it takes GLIPH to process just 500. More importantly, irrespective of CPU or GPU hardware, TCR-BERT exhibits linear runtime scaling, making it much more scalable for analyzing large datasets.

**Supplementary Figure 7:**
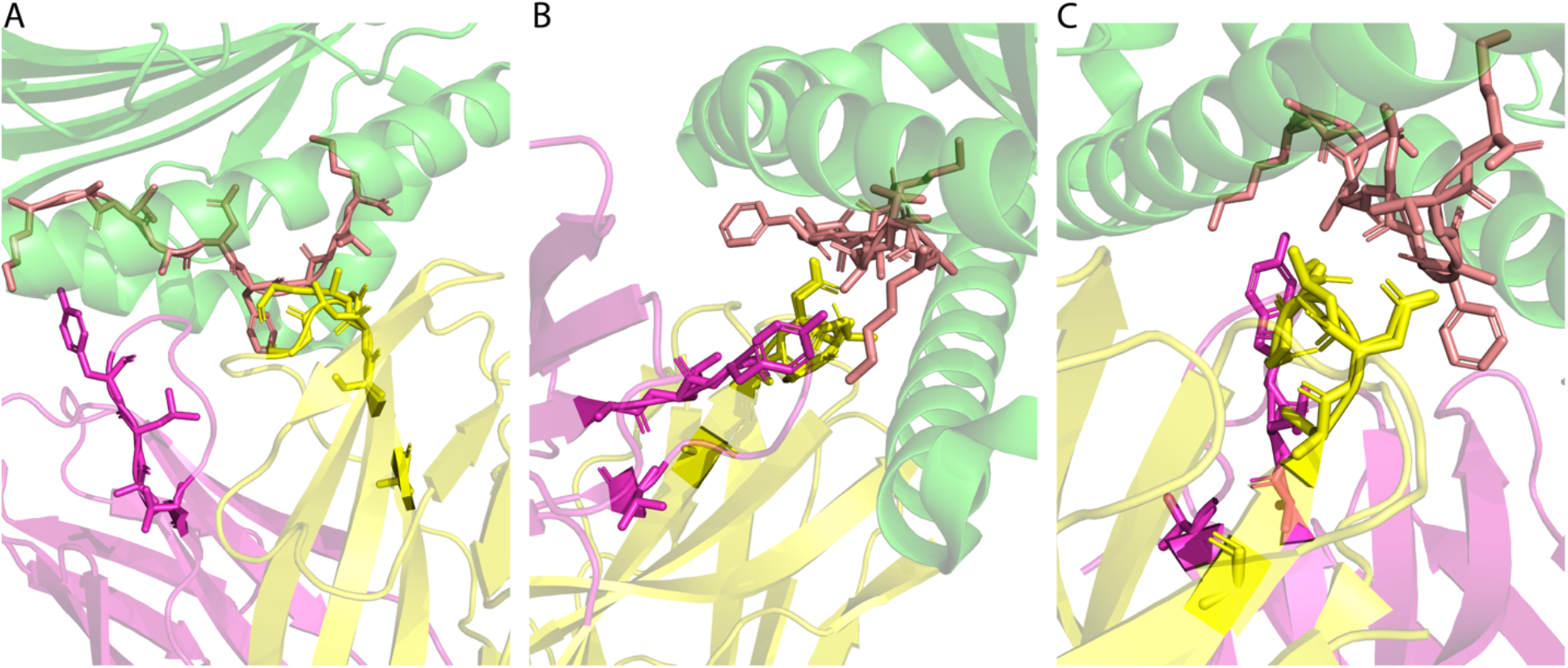
Additional views of 5m00 with high-attention TRA/TRB residues highlighted. 3D structures showing the MHC (green), modified GP33 antigen (salmon), and TRA (pink) and TRB (yellow). GP33 antigen side chains are shown, as well as side chains for TRA and TRB residues with TCR-BERT attentions in the top 33^rd^ percentile of attention values in each chain. Other residues are shown in faded cartoon-ribbon illustrations without side chains. Attention values are derived from average attentions across test set sequences with identical lengths as sequences profiled in PDB structure 5m00. In all views, the TRA/TRB residues with the greatest attention are frequently in direct contact with the antigen peptide’s side chains. (A) Flipped view of Figure 4D (B) View with the TRA in the foreground (C) View with the TRB in the foreground

**Supplementary Figure 8:**
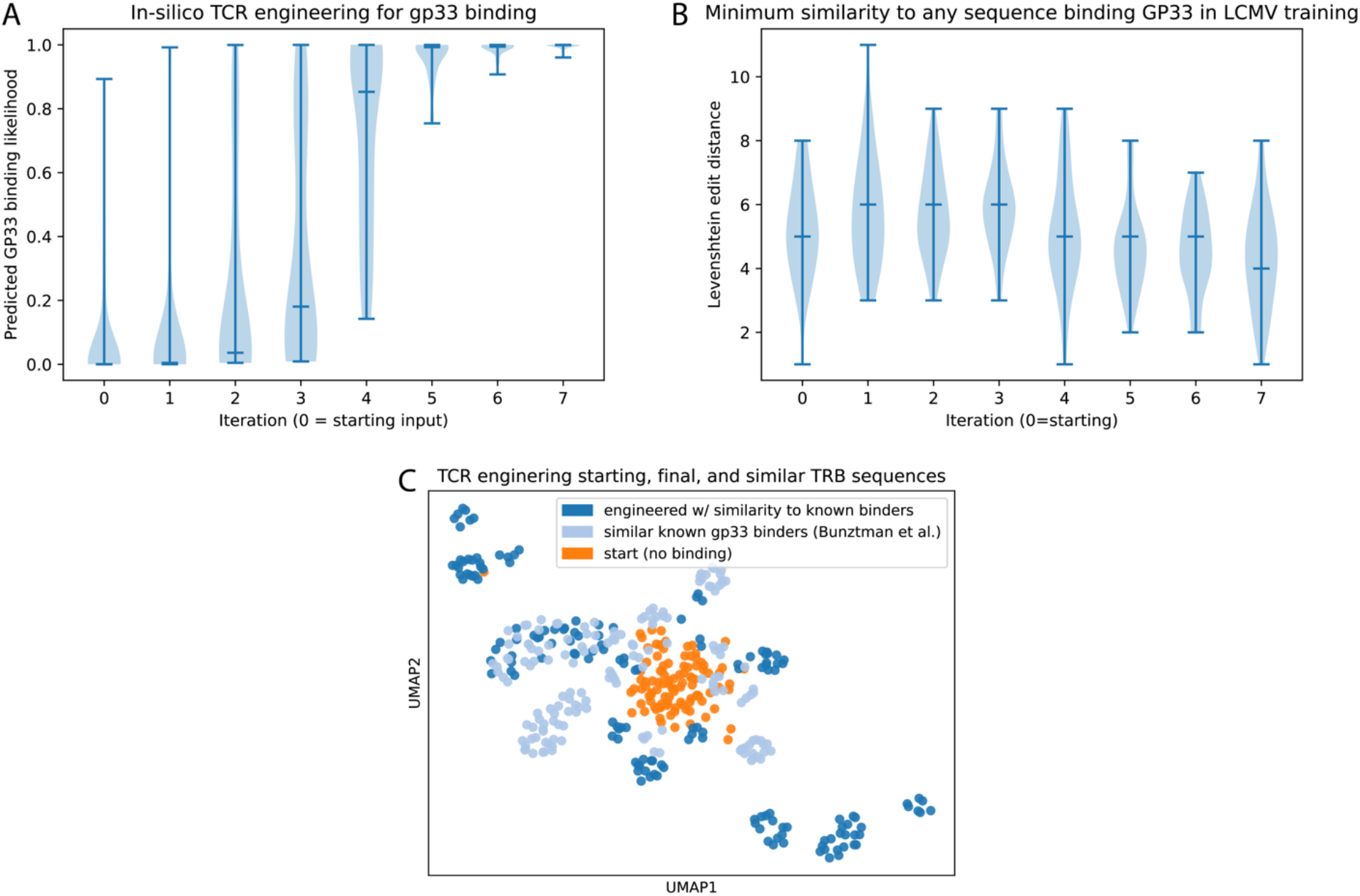
TCR engineering supplement. (A) Our algorithm for engineering TCR sequences successfully increases the predicted binding of sequences (y-axis) with each iteration (x-axis). The 0-th iteration represents the input sequences. We stop our engineering iterations (Figure 5A) when all sequences exceed 95% predicted binding probability, which occurs after 7 iterations in this instance. (B) Our algorithm for engineering TCR sequences for antigen binding works without simply regurgitating training sequences. We show this by computing the Levenshtein edit distance (larger values indicate greater dissimilarity, y-axis) between each generated TRA/TRB sequence pair and the most similar sequence pair in our dataset of GP33-binding TCRs. This is expressed as a violin plot for each iteration of our procedure (x-axis). At no point during the TCR engineering process do we include any sequence directly seen in training (i.e., no instances of 0 edit distance). This indicates that we are generating truly novel sequences that TCR-BERT has not seen before. (C) We can also visualize the progression from starting to engineered sequences in TCR-BERT’s embedding space. Here, we embed each TRB sequence (using the same variant of TCR-BERT pre-trained on MAA and antigen classification as we use for other embedding visualizations for consistency) and visualize the embeddings using UMAP. Colors correspond to starting, engineered and similar binders (i.e., TRBs identified in a separate experiment to also bind GP33 that bear significant similarity to our final engineered set). Known GP33 binders (light blue) lie towards the “outskirts” of our starting set (orange), and our engineered set pull outwards toward these (dark blue). This shows that our TCR engineering process explores the landscape of TCR sequences without being confined to the area spanned by input sequences.

**Supplementary Figure 9:**
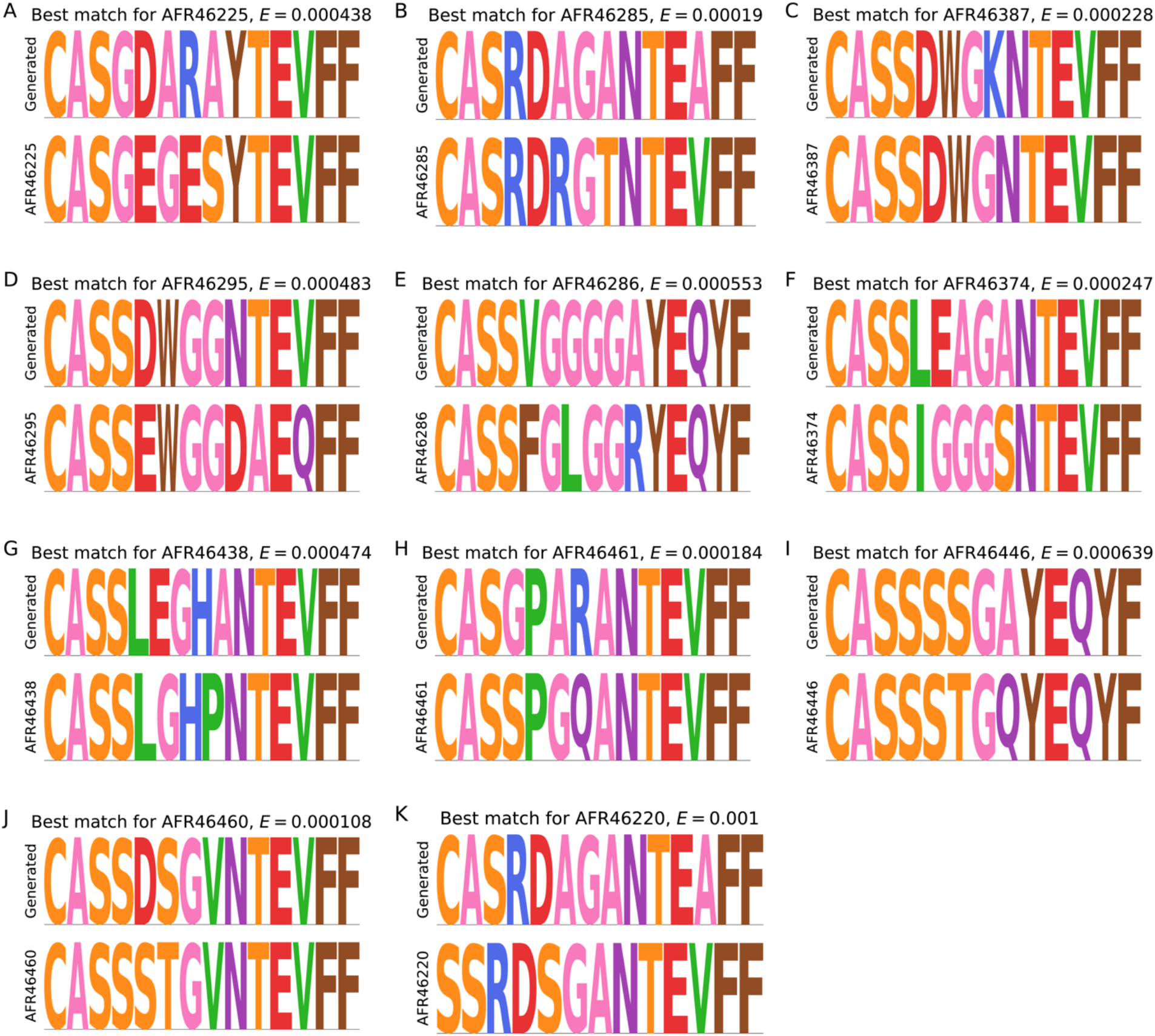
Detailed sequence comparisons for 11 novel matches. Each of these 11 matches was not present in the initial pool of matches corresponding to the starting set of sequences for TCR engineering. For each match (A-K, bottom panels), we show the generated TRB with the highest bit score/lowest E-value to that match (A-K, top panels). Each pairing is annotated with the E-value corresponding to that match.

**Supplementary Figure 10:**
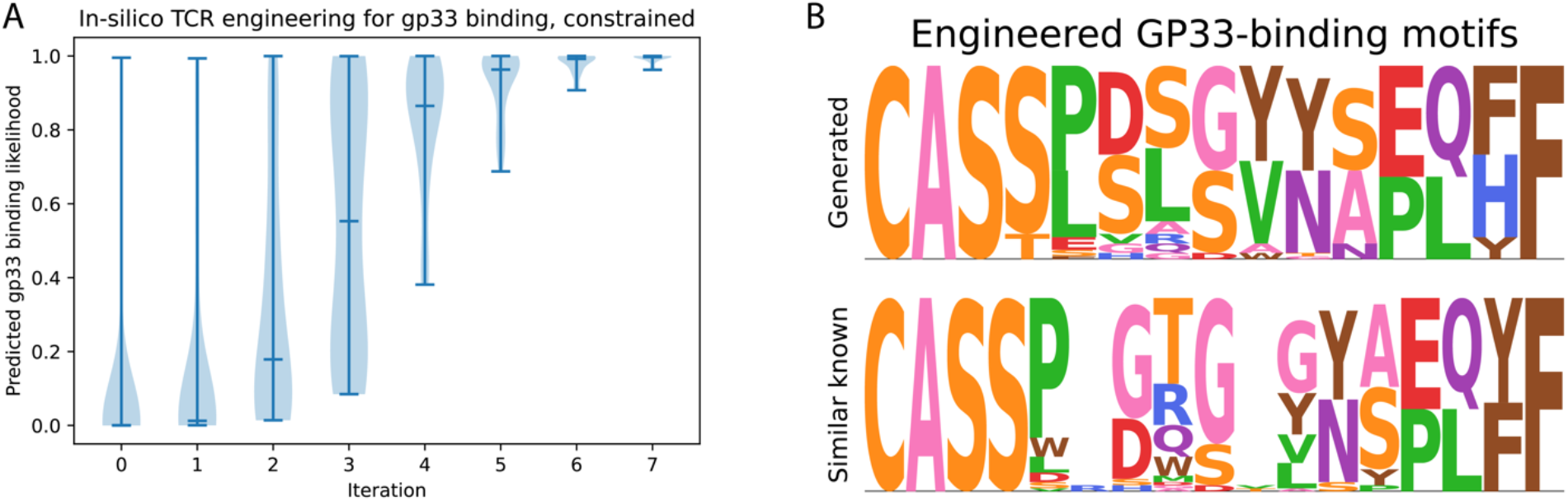
Additional TCR engineering results. (A) We repeat TCR engineering with a different starting set of negative sequences, also drawn from the pool of LCMV test set negatives. As before, our TCR engineering procedure successfully creates sequences with increasingly greater predicted binding with each iteration. (B) We match the final set of engineered TRBs to known murine TRBs using BLAST (E-value ≤ 0.001). Among the significant hits, 61/189 correspond to GP33-binding TRB sequences. The bottom motif corresponds to these hits, and the top motif corresponds to our corresponding generated sequences. For context, proportionally fewer (69/393) matches for the starting input set corresponded to GP33-binding TRBs. As before, TCR-BERT proportionally enriches for GP33 binders.

